# Alcohol drinking is associated with greater calcium activity in mouse central amygdala dynorphin-expressing neurons

**DOI:** 10.1101/2024.02.18.580880

**Authors:** Christina L. Lebonville, Jennifer A. Rinker, Krysten O’Hara, Christopher S. McMahan, Michaela Hoffman, Howard C. Becker, Patrick J. Mulholland

## Abstract

Alcohol use disorder (AUD) is a chronic disease that poses significant economic burden and health risks. It is pivotal to better understand brain mechanisms engaged by alcohol that promote misuse. The central amygdala (CeA) has emerged as a key mediator of excessive preclinical alcohol consumption. A dynorphin-expressing subpopulation within the CeA (CeA^Dyn^) has been implicated in excessive alcohol drinking, yet how cellular activity of CeA^Dyn^ neurons relates to ongoing alcohol drinking is not well-understood. The current study interrogated the engagement of CeA^Dyn^ neurons in male and female mice during voluntary alcohol consumption using fiber photometry and compared this cellular response with that of other solutions having similar motivational and/or taste characteristics. Activity of a calcium sensor, GCaMP7f, expressed in mouse CeA^Dyn^ neurons was recorded and time-locked to bouts of drinking. Multilevel linear mixed modeling was applied to better resolve focal effects from complex data. These analyses revealed a relatively large increase in CeA^Dyn^ neuron calcium transients after bouts of alcohol drinking compared to water or sucrose drinking, indicating these neurons are uniquely engaged during alcohol consumption. Drinking behavior unique to alcohol (i.e., longer bout durations) did not fully explain signal differences between alcohol and other solutions nor did the relatively increased alcohol response diminish over time. No other conditions or solutions tested reproduced the pronounced change in CeA^Dyn^ activity associated with alcohol drinking. These findings, collectively, support the presence of a unique functional signature for alcohol in a cell population known to control excessive alcohol drinking.

**Highlights:**

- Central amygdala dynorphin cells (CeA^Dyn^) are firmly implicated in alcohol misuse.
- CeA^Dyn^ neuron activity was higher when mice drank alcohol versus other solutions.
- Neither how mice drank alcohol nor motivational states could explain this activity.
- CeA^Dyn^ neurons having uniquely high alcohol responses may underlie AUD development.

## Introduction

Alcohol use disorder (AUD) is a prevalent chronic relapsing disease, affecting 30 million people (~9%) in the United States alone (SAMHSA, 2022) and poses significant negative economic and health consequences (Sacks et al., 2015; WHO, 2018). Evidence from human and animal studies has revealed brain regions involved in excessive alcohol (ethanol) drinking and the development of AUD, including the amygdala (for review see: Roberto et al. 2020). In rodent models of excessive alcohol drinking, the central amygdala (CeA) has emerged as a key mediator of alcohol consumption, particularly in alcohol dependence. The CeA has three sub-nuclei comprised of heterogeneous GABAergic neurons that express different neuropeptides. A dynorphin-expressing subpopulation within the CeA (CeA^Dyn^), like other CeA neuropeptide subpopulations, has been implicated in excessive alcohol drinking across both acute and chronic alcohol exposure models (for review see: Walker et al. 2021). CeA^Dyn^ are known to regulate both binge alcohol drinking and stress-enhanced drinking in alcohol dependence (Anderson et al., 2019; Bloodgood et al., 2020; Haun et al., 2022). Dynorphin signaling disruption within the CeA, by either chemogenetic inhibition, receptor antagonism, or conditional knockout, decreases alcohol drinking. Conditional knockout of the CeA dynorphin precursor, prodynorphin (*Pdyn),* also decreased drinking in a sex-dependent manner and, similarly, prolonged alcohol exposure increases excitability of CeA^Dyn^ neurons in male, but not female mice (Bloodgood et al., 2020). While CeA^Dyn^ neuronal activity was previously measured in both sexes during alcohol drinking (Roland et al., 2024), these studies were conducted under water-deprived conditions with brief momentary access (≤10 min), sometimes interleaved with access to other solutions in the same session, making it challenging to relate sex-specific CeA^Dyn^ activity to alcohol drinking during typical alcohol exposure paradigms. The current studies were designed, therefore, to investigate CeA^Dyn^ neuron activity dynamics during a limited alcohol access paradigm without water deprivation to better understand alcohol-associated CeA^Dyn^ neuron responses across sex.

To capture real-time activity of CeA^Dyn^ neurons during voluntary consumption of alcohol or other solutions, we used in vivo fiber photometry coupled with lickometry circuits in freely behaving male and female *Pdyn*-IRES-Cre mice. While conducting these experiments, we encountered two known challenges to the fidelity of such data – control photometry signal artifacts and inaccurate lickometry circuit triggers – and developed tested methods to overcome these challenges. Additionally, because of the complex structure of repeated measures data using a within-subjects experimental design, like that acquired in this fiber photometry drinking study, we applied a multilevel linear mixed model (MLMM) statistical approach for notable advantages over standard photometry analyses (see (Gueorguieva and Krystal, 2004) for comparison). Specifically, MLMMs are designed to account for repeated measures data even with missing data or unbalanced designs (West et al., 2022). Compared to repeated measures or nested ANOVAs that consider random variation from a single source (usually within subject), MLMMs can better estimate composite effects of multiple random variables from a complex, nested data structure, providing more power to detect and accurately estimate effects from critical variables (see (Aarts et al., 2014)). This is relevant to the current study in that mice showed individual variation within and across drinking sessions and solutions. Linear mixed modeling, more generally, has been proposed as a more powerful alternative to standard photometry analysis methods (Simpson et al., 2024) and has been previously applied to photometry data (Cazares et al., 2022). Applied here, MLMMs provided a statistically sound foundation to test both primary and secondary hypotheses on the relationship between alcohol drinking and CeA^Dyn^ neuron calcium activity. We hypothesized that CeA^Dyn^ neurons would show distinct signaling around a bout of alcohol drinking compared to other substances, given the substantial evidence of this population’s importance to alcohol consumption and that males would show a more pronounced pattern of activity compared to females, due to reported sex differences in the engagement of CeA^Dyn^ neurons after chronic alcohol. Collectively, these studies aimed to understand early alcohol engagement of CeA^Dyn^ neurons in both sexes to better understand the role of these responses in promoting excessive alcohol drinking.

## Methods

### Animals

Male and female prodynorphin-Cre (*Pdyn-*IRES-Cre) mice (n = 35, 17 male, 18 female) were bred in-house from homozygous *Pdyn*-IRES-Cre mice (expressing Cre-recombinase under the prodynorphin promoter, from which dynorphin is cleaved; B6;129S-Pdyn^tm1.1(cre)Mjkr^/LowlJ, JAX Stock #027958) and C57BL/6J mice (JAX Stock #000664). Mice were 8-17 weeks of age at the start of the experiments and allowed *ad libitum* access to food throughout and water outside of experimental 2-hr drinking sessions. Mice of both sexes were individually housed in a single colony room on a reversed 12-hr light-dark cycle (lights off at 9 am). All animal procedures were approved and facilities inspected by the Medical University of South Carolina’s (MUSC) Institutional Animal Care and Use Committee (IACUC) in accordance with the guidelines established by the National Research Council (“The Guide for the Care and Use of Laboratory Animals,” 2016). MUSC is accredited by the Association for the Assessment and Accreditation of Laboratory Animal Care (AAALAC) and meets U.S. Department of Agriculture and Public Health Service standards and regulations. Mice for Experiment 1 were run in two independent, mixed-sex cohorts and for Experiment 2 run in a single mixed-sex cohort. Mice that did not show spontaneous acute GCaMP7f activity were removed at the onset of the study. Mice determined to have off target ferrule or viral placement were removed from the final analysis after histological verification.

### Surgery & Viruses

Mice underwent unilateral surgical delivery of a Cre-dependent GCaMP7f calcium sensor into the CeA (pGP-AAV1-syn-FLEX-jGCaMP7f-WPRE, a gift from Douglas Kim & GENIE Project, Addgene viral prep # 104492-AAV1; RRID: Addgene_104492, Lots v49556 and v118530 with titers of 1-2 x10^13^ GC/mL) and implantation of a fiber optic ferrule (MFC_400/430-0.57(or 0.66)_5.3mm_ZF2.5(G)_FLT, Doric Lenses Inc, Quebec, QC, Canada) above the target CeA population. The virus was delivered at a volume of 200 nL and rate of 100 nL/min. Following infusion, injectors remained in place for 10 min to allow for diffusion away from the injector tip before injector retraction. Target stereotaxic coordinates for CeA were −1.4 AP, +2.7 ML, −4.7 virus/−4.62 fiber DV (in mm relative to bregma) based on the mouse stereotaxic atlas (Franklin and Paxinos, 2008). Mice were administered carprofen (5 mg/kg, SC), xylocaine (2%, topical), and bacitracin/polymixin (2%, topical) at the time of surgery and for at least 2 days post-surgery. Mice were then given at least 5 weeks to recover from surgery and for viral incubation before being acclimated to tethering procedures and tested for GCaMP7f signal.

### Limited Access to Alcohol and Other Solutions

Mice were given consecutive access to a single bottle of test solution 3 hours into the dark cycle in their home cages for 2 hours/day, 5-6 days/week. Identical graduated sipper tubes were used for all solutions and access was given to all mice, even though photometry was recorded from two mice each day. For Experiment 1, mice were first given access to 20% (v/v) alcohol for 3 weeks, water for two weeks, and then 0.5% sucrose (w/v) for three weeks. For Experiment 2, mice had sequential access to multiple quinine concentrations, water, water following ~24-hr water deprivation, 0.5% (w/v) sucrose plus low quinine concentration, and 0.5% (w/v) sucrose plus high quinine concentration. For the first week of quinine access, we started with a 30 mM concentration of quinine and reduced the concentration daily until most mice began to consume the solution. This approach was chosen to allow capture of neural activity at the most aversive quinine concentration voluntarily consumed. The first concentration to be consumed by most mice (125 μM, referred to as a “high” concentration) was presented daily the following week. Since 64.5% of the bouts for the high quinine concentration came from two mice, the concentration was reduced to 90 μM (referred to as a “low” concentration) for 2 weeks of access. For each subsequent solution, access was given for 1 week. Solutions were made from commercially available chemical solids (sucrose, S5-3, ThermoFisher Scientific; quinine hydrochloride dihydrate (QHCl), Q1125, MilliporeSigma) or high purity stock solutions (95% ethanol (EtOH), 2801, Decon Labs). Solutions other than alcohol and water were stored long-term at 4°C protected from light but were allowed to warm to room temperature before drinking sessions. Licking behavior was identified as a bout if lick frequency was >4.5 Hz, interlick interval was <500 ms, and duration was >500 ms based on the known microstructure of rodent drinking behavior (Boughter et al., 2007; Davis and Smith, 1992; Johnson et al., 2010). Sipper contacts were recorded and time-locked to photometry data using lickometer circuits (Med Associates Inc., Fairfax, VT, USA). Overhead video recordings of drinking sessions were collected, and a subset were initially used to confirm that licking bouts were truly consummatory and not any other interaction with the spout (e.g., sniffing, licking the spout side rather than the opening, touching with paws). Validation of each bout was critical for our analytical approach since every data point is used to build the MLMM whereas in more classical approaches, the influence of a small amount of erroneous data may have less influence. To validate licking bouts, therefore, we trained a deep-learning model using DeepEthogram (Bohnslav et al., 2021) to distinguish consummatory licks from all other sipper contacts. The final applied model had 95% average individual frame classification accuracy and >98% bout classification accuracy when compared to manual evaluation. When the trained DeepEthogram model was applied to experimental data, 9-11% of all bouts were identified as invalid and removed from the final analysis. Within high concentration quinine sessions (≥125 µM), 30-67% of bouts were identified as non-consummatory spout contacts, highlighting the importance of bout verification, particularly during access to potentially aversive solutions.

### Fiber Photometry Recording and Processing

Fiber optic patch cords containing rotary joints (Doric; 0.57 NA; 400 μm) were mounted within custom-built balance arms to allow free exploration and head movement. Patch cord terminals were connected to sterilized chronic indwelling optic fiber cannula (Doric; 0.57-0.66 NA) using pinch connectors (Thorlabs Inc., Newton, NJ, USA). After habituating mice to tethering procedures, CeA^Dyn^ neuronal calcium transients were recorded from two mice during each drinking session. Consecutive recording sessions were limited to two days from an individual mouse to avoid GCaMP7f photobleaching. During recording, GCaMP7f was excited using a collimated 470 nm LED (Thorlabs), and signal from a concurrent, equally powered 405 nm LED was used for normalization to control for movement artifacts. Illumination power for each LED (5-30 µW) was sinusoidally modulated at 531 Hz (405 nm) and 211 Hz (470 nm) and was passed through Doric fluorescent Mini Cubes. Emission light was focused onto a femtowatt photoreceiver (Newport, model 1672151; DC low setting) and sampled at 6.1 kHz by a RZ5P lock-in digital processor controlled by photometry processing software (Synapse, Tucker-Davis Technologies, Alachua, FL, USA). The GCaMP7f signal was integrated in real-time with licking behavior using lickometer circuits (Med Associates Inc., Fairfax, VT, USA) following our previous methods (Rinker et al., 2023). Fiber optic patch cords were photobleached twice weekly overnight using a 405 nm LED to reduce autofluorescence. Although the magnitude of photobleaching across the session was small, our initial analysis revealed significant differences in the slope of the 405 and 470 nm signals over time (slope, median 405 nm = −5.06×10^−3^; 470 nm = −3.41×10^−3^; n = 127/channel; Wilcoxon Signed Rank test, V = 7556, p < 0.0001, two-sided). This is a known phenomenon from uneven photobleaching between the two wavelengths (Simpson et al., 2024). Thus, a linear least-squares fit was applied to detrend each channel separately before further pre-processing. Raw, demodulated emission signals were detrended, normalized, and processed using custom MATLAB scripts according to our standard protocol (Rinker et al., 2023) with the following modification.

We, like others, observed inverted 405 nm signal during periods of increased 470 nm signal ((Barnett et al., 2017; Simpson et al., 2024); see **Figure 1A**). To eliminate these calcium-dependent control fluctuations, we applied a correction algorithm and investigated the ability of this strategy to reduce the influence of these deflections on the final, normalized percent delta*F*/*F (%*Δ*F/F)* signal. We developed a Jointly Constrained B-spline model (JCBM) using machine learning that directly identifies and removes the sustained deflections in the 405 nm signal (see *Machine Learning Pipeline for 405 nm Correction*). We then compared the detrended, unnormalized 470 nm signal (“470 UN”), standard normalized %Δ*F*/*F* signal using an uncorrected 405 nm signal (“Standard”), normalized %Δ*F*/*F* signal generated using a robust linear modeling algorithm (“Robust”) following previous methods (Martianova et al., 2019), and normalized %Δ*F*/*F* signal using the new JCBM method. To evaluate the performance of each pre-processing strategy, we separately analyzed signals during large 405 nm signal deflections, measured large-scale signal dynamics, and analyzed signals during movement artifacts. For the first analysis, we measured the signal generated using each pre-processing method during periods of large, sustained deflections in the 405 nm signal. A MATLAB script was used to automatically identify these periods within Experiment 1 data and extract identical segments in data from each normalization method. After visually curating the longest and most stable deflections, the final data for this analysis contained 392 events across session, mouse, sex, and solution. To compare how each normalization method influenced resulting signal dynamics across an entire 2-hr recording, we analyzed signal mean absolute deviation (MAD) and the mean amplitude and frequency of identified signal peaks in 135 sessions across sex and solution from Experiment 1 data. For the final analysis, we artificially generated 27 movement artifacts across Experiment 2 mice after the conclusion of testing by hand restraining them and rapidly flipping them upside down around the body’s long axis. By marking precisely when these movement artifacts were generated, we were able to isolate signal deflections due to movement and analyze the ability of each normalization method to remove these artifacts. Based on these analyses, shown in *Results*, we determined that the JCBM method produced the least distorted normalized signal and this method was therefore used for all subsequent analyses. Briefly, after detrending each channel separately, %Δ*F*/*F* = [(470 nm signal – JCBM-corrected 405 nm signal)/JCBM-corrected 405 nm signal]*100. We extracted mean signal 1 sec before and after a drinking bout and standardized these values by subtracting the mouse-specific experiment-wide mean and dividing by the mouse-specific experiment-wide standard deviation of these signals to produce a within-mouse z-score (z%Δ*F*/*F*). Our primary dependent variable was the change (slope) between standardized before and after bout GCaMP7f signals as this was used to measure bout-specific activity. In the absence of a change in bout-specific activity, the average of standardized signal before and after licking bouts was used to assess general shifts (intercepts) in GCaMP7f activity.

**FIGURE 1.**
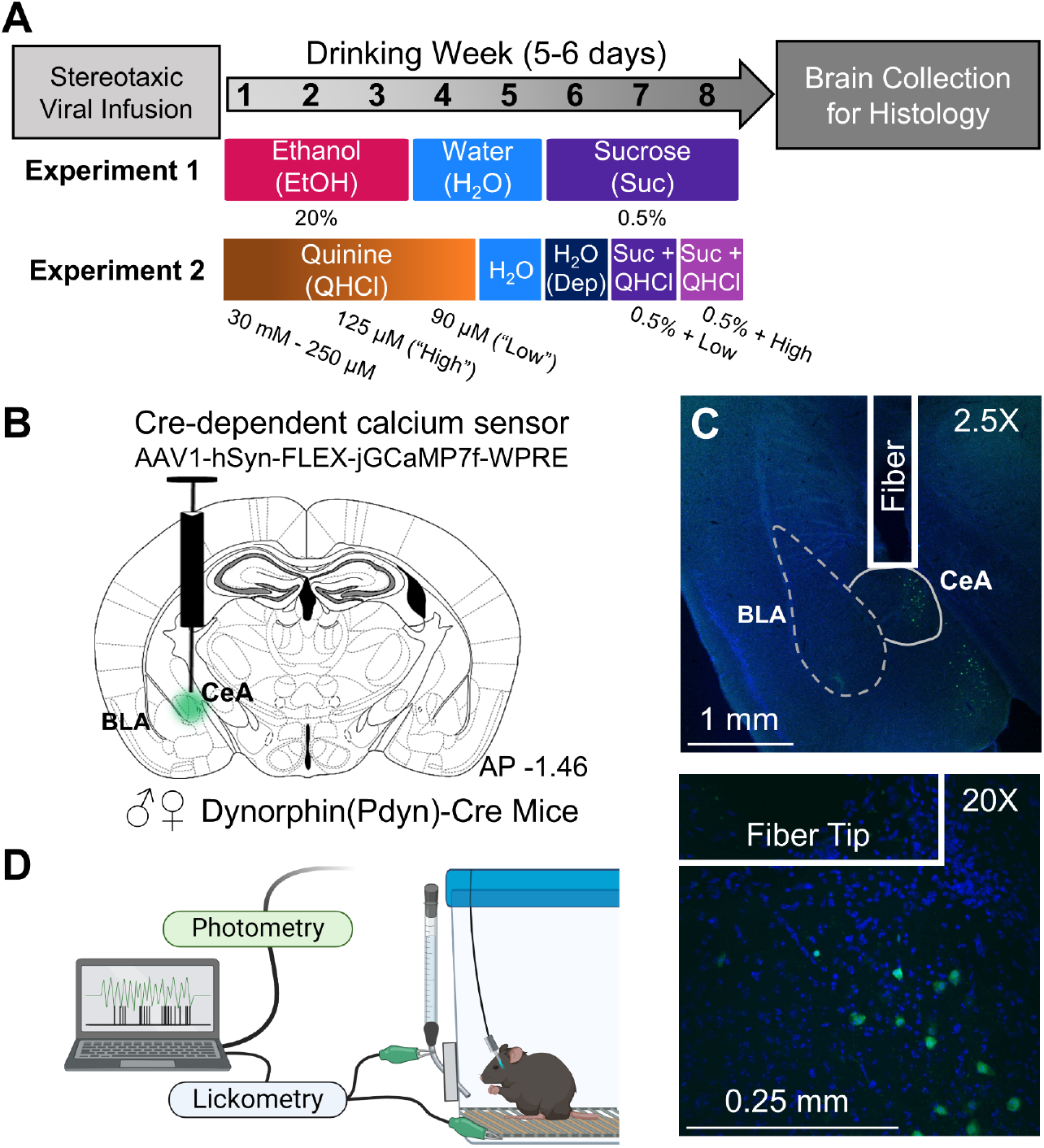
(**A**) Experimental design and timeline for Experiments 1 and 2. (**B**) Strategy for viral infusion of Cre-dependent GCaMP7f into the CeA and fiber optic ferrule implantation above injection site to record calcium transients from CeA^Dyn^ neurons. (**C**) Representative images of viral expression and fiber optic ferrule placement at 2.5X (top) and 20X (bottom) magnification. (**D**) Lickometry circuit used to integrate home cage drinking with fiber photometry recording of CeA^Dyn^ activity. Created in BioRender. Mulholland, P. (2024) https://BioRender.com/w82c805

### Machine Learning Pipeline for 405 nm Correction

The goal of our novel machine learning pipeline was to correct Ca^2+^-dependent downward deflections in the 405 nm signal during periods of high signal in the 470 nm channel. To this end, we conceptualized a model for the 470 nm series, 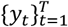, and the 405 nm series, 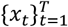, that is given by

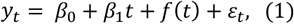

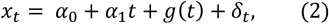

where *β*_0_ (*α*_0_) is the usual intercept, *β*_1_(*α*_1_) is a slope parameter that captures the general trend of the series, *f* (*g*) is a time dependent function that captures the fluctuation around the general trend, and *ε*_*t*_ (*δ*_*t*_) is a random error term. To provide a flexible specification, we make use of B-splines to model *f* and *g*; i.e., we specify

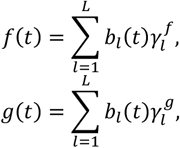

where 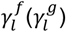 are spline coefficients that are unique to *f* (*g*) and *b*_*l*_ (*t*) are B-spline basis functions which are fully determined once a knot set and degree are specified. In our implementation, we made use of a knot set consisting of 147 interior knots placed at equally spaced quantiles of the series and specified the degree to be 3, yielding 150 basis functions; for further discussion on B-splines and their uses, see (Boor, 1972). Our use of B-splines in this context afforded us two primary benefits. First, our specifications allowed for a great deal of modeling flexibility rendering our model more than adept at capturing fluctuations around the trend. This flexibility, however, came at the expense of potential overfitting issues. To mitigate this potential, we regularized the estimation of the spline coefficients via the following penalty

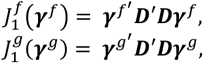

where 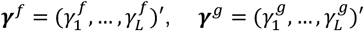and ***D*** is a (*L* − 1) × *L* − dimensional first order difference matrix; i.e., ***D***_*ll*_ = 1 and ***D***_*l,l*+1_ = −1, for *l* = 1, …, *L* − 1, and ***D***_*ll*_ = 0 otherwise, for further discussion see (Eilers and Marx, 1996). The second benefit of this specification, is it afforded us the ability to enforce functional similarity between *f* and *g* through the following penalty

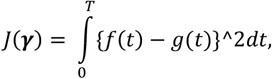

where ***γ*** = (***γ***^*f*^, ***γ***^*g*^)′. Leveraging the spline representation, this penalty can be expressed in matrix form as follows *J*(***γ***) = ***γ***^′^***J γ***, where

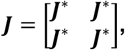

and *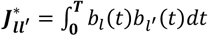*. Finally, the last regularization term that we adopted is a ridge penalty that acts to shrink regression coefficients toward 0; i.e., we specified the following penalty

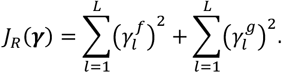

Thus, the objective function used to fit (1) and (2) is given by

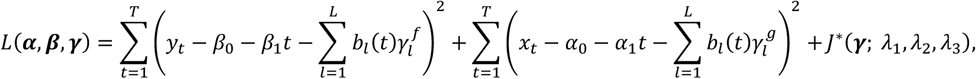

where 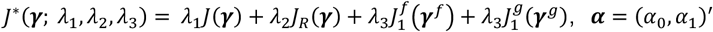, and ***β*** = (*β*_0_, *β*_1_)^′^. A few comments are warranted. First, the minimizer of the objective function can be computed in closed form for any value of the penalty parameters *λ*_1_, *λ*_2_, and *λ*_3._ Second, different configurations of the penalty parameters ultimately yield different model fits. To select the best set, we utilized the BIC criterion with the degrees of freedom being computed as the trace of the smoothing matrix (for further discussion of this strategy see (Hastie et al., 2009)). Given the parameter estimates arising from this process, we made use of the fitted values to correct the detrended 405 nm signal before 470 nm normalization.

### Histological Validation & Final Dataset

After behavioral experiments, mice were sacrificed by injection with urethane (1.5 g/kg, 50% w/v, IP) and, at surgical anesthetic depth, transcardially perfused with ice-cold 1X Dulbecco’s phosphate buffered saline (PBS) and 4% paraformaldehyde solutions. Brains were extracted and stored in 4% paraformaldehyde solution at 4°C until further processing. Brains were sliced into 40 µm sections on a vibratome (VT100S, Leica Microsystems, Wetzlar, Germany) which were then slide mounted to charged glass slides and coverslipped using VECTASHIELD Vibrance® with DAPI (Vector Labs, Burlingame, CA, USA) or ProLong™ Glass with NucBlue™ (ThermoFisher Scientific, Waltham, MA, USA). On-target and dorsal placement of the fiber ferrule above GCaMP7f expression was visually verified using a fluorescent microscope (EVOS fl, AMG, Bothell, Washington, USA). Representative images were captured using a confocal microscope (Carl Zeiss LSM 880, Oberkochen, Germany). For Experiment 1 (**Figure 1A**), of the 26 (13 female/13 male) *Pdyn*-IRES-Cre mice who underwent surgery to express GCaMP7f in the CeA, one female and one male mouse were lost due to surgical complications and six female and two male mice were removed early from the experiments due to having low spontaneous calcium signal. Following experimentation, we histologically validated GCaMP7f expression and fiber placement within the CeA (**Figure 1C)** and removed two males for nonspecific virus expression. Two additional bouts were excluded from analyses because their bout characteristics were far outlying visually and statistically (Grubbs). The final dataset included calcium and drinking data from n = 14 mice (8 male, 6 female) during 122 sessions and 2832 validated drinking bouts. For Experiment 2 (**Figure 1A**), of the 9 (5 female/4 male) *Pdyn*-IRES-Cre mice who underwent surgery to express GCaMP7f in the CeA, one male mouse was lost due to surgical complications. All mice had histologically validated GCaMP7f expression and fiber placement within the CeA. The final dataset included calcium and drinking data from n = 8 mice (3 male, 5 female) during 52 sessions and 1243 validated drinking bouts. This dataset can be accessed at [*data repository link to be added at publication*].

### Statistical Analysis and Multilevel Linear Mixed Modelling

Linear mixed models (including hierarchical or multilevel) contain both fixed (independent) factors that test effects of interest and random factors whose effects are estimated and used to account for sources of variability other than fixed factors. This approach gives better resolution of fixed effects but determining appropriate random factors requires careful consideration. We empirically tested various random effects structures for each outcome (dependent) variable using an unconditional means model, absent any fixed factors. We selected random effects structures from tested candidates that were logical given the data, were appropriately levelled for the outcome, gave the highest ICC and lowest AIC/BIC (metrics of clustering within random effects and model fit/parsimony), and whose omission significantly changed model fit determined by a Log Likelihood Test. Random effects structures for the main analyses (i.e., calcium signal relating to drinking behavior) are listed in **Table 1**. For models in Experiment 1, we used Helmert contrasts for categorical variables (i.e. Sex, Time, Solution). A Helmert contrast for the three-level Solution factor tested the primary hypothesis that alcohol would differ from both other solutions and gave two comparisons: sucrose compared to water and then alcohol compared to water and sucrose, jointly. A Helmert contrast also provided the benefit of centeredness and orthogonality, thus reduced problems with variance estimation (Yaremych et al., 2021), maximized statistical power. As we had more specific hypotheses for normalization method and Experiment 2 analyses, we used effect coding with the primary factors of interest (i.e., Method and Solution, respectively) to compare each condition to a control condition (470 UN or water). To avoid estimation bias in upper-level interaction effects and to increase interpretability, scaled mean-centered forms of continuous independent variables were used. Levels of a variable described as “high” or “low” refers to estimates at +1 or –1 standard deviation from the mean, respectively. The variable “session” was coded as the day of drinking within each solution. All tests other than model comparisons (which used Maximum Likelihood) were performed using Restricted Maximum Likelihood (REML) estimation in R (R_Core_Team, 2023) with custom scripts (available in GitHub repository to be made available in final article), the “lme4” package (Bates et al., 2015), and guidance from (West et al., 2022). Post hoc tests with Tukey adjustment for multiple comparisons and estimated marginal means were computed using “emmeans” and “stats” R packages (Lenth, 2023; R_Core_Team, 2023). Final models were evaluated for multicollinearity using generalized variance-inflation factors (GVIFs) from the “car” R package (Fox and Weisberg, 2018) where GVIFs < 5 were considered to have no issues. Degrees of freedom for t-statistics were estimated with the R package “lmerTest” (Kuznetsova et al., 2017) using Satterthwaite approximations, which produce acceptable Type I error rates (Luke, 2017). Data were visualized using R package “ggplot2” (Wickham, 2016) with numerous add-on packages (system environment available in *Supplement A* for reproducibility).

**TABLE 1.** Multilevel linear mixed model (MLMM) design used in Experiments 1 and 2.

| Level | Fixed Variables<br>(Bold = Primary Interests) | Random Variables | Outcome Variables |
| --- | --- | --- | --- |
| Mouse (Lvl 4) | **Time** (Lvl 1), **Solution** (Lvl 3), **Sex** | [Exp 1: Time + Solution (slopes) or Exp 2: Time (slope)] / Session nested within Mouse (intercept)] + [Bout nested in Session within Mouse (intercept)] + [Bout (intercept)] | Signal |
| Session (Lvl 3) | **Time** (Lvl 1), **Solution** (EtOH, Water, Sucrose), Session-Level Characteristics: Consumption (mL/kg) | [Mouse (intercept)] | Session-Level Drinking Characteristics |
| Bout (Lvl 2) | **Time** (Lvl 1), Bout-Level Characteristics: Bout Duration, Onset Time (Position) | [Session nested within Mouse (intercept)] | Bout-Level Drinking Characteristics |
| Time Around Bout (Lvl 1) | **Time** (Before, After) |  |  |

For analyses of normalization method effects on signal during 405 nm deflections we used an MLMM with Method (470 UN, Standard, Robust, and JCBM), Sex (Male, Female), and Solution (EtOH, Water, Sucrose) as fixed factors and Mouse as a random intercept. For analyses of normalization method effects on signal dynamics across the entire 2-hr session, we used MLMMs with Method (470 UN, Standard, Robust, and JCBM) and Solution (EtOH, Water, Sucrose) as fixed factors and Mouse and Session nested within Mouse as random intercepts. Sex as a fixed factor was not supported for these outcome variables since a Log Likelihood Test comparing the fit of the models with and without Sex was non-significant (χ^2^_4_ = 3.123-4.729, *ps* ≥ 0.3162) and adding Sex as a factor did not result in any Sex effects nor interactions. Solution was excluded from the peak frequency model as it did not significantly improve model fit (Log Likelihood Test χ^2^_8_ = 8.064, *p* = 0.4272) nor add any effects or interactions. For analyses of normalization method effects on correcting movement artifacts, we used a MLMM with Method and Sex as fixed factors for the slope outcome variable. There was a significant interaction between Sex and Method, but for clarity, slope is presented collapsed by Sex. Specifically, males showed smaller movement artifact normalization in 470 nm slope than females for two of the three methods, but this was likely due to larger pre-normalization movement artifacts in females, rather than normalization method *per se*. We used Method alone for the correlation outcome variable as Sex did not significantly improve model fit (Log Likelihood Test χ^2^_4_ = 8.781, *p*= 0.0668) nor add any effects or interactions for this variable. Mouse was used as a random factor (intercept) for both. Mouse as a random factor did not significantly change the model fit for the slope model, but we decided to include it to be conservative about the repeated measures within mouse. Full factorial statistics can be found in *Supplement B*.

Analyses of the relationship between CeA^Dyn^ calcium activity and drinking behavior first fit the data with a best-fit MLMM and then interpreted parameter estimates of the effects. Our drinking characteristic (i.e., consumption, bout duration) MLMM models included Sex and Solution as fixed variables and outcome-optimized random effects (**Table 1**). Our signal MLMM model included fixed factors Sex, Solution, and Time around bout. Based on random effect optimization for signal models, we allowed the intercept to vary for each session within mouse, each bout within session and mouse, and each bout unnested due to bout being also crossed between mice and solutions (**Table 1**). Since we had sufficient power with this outcome variable for Experiment 1, we also included random slopes for Time and Solution with the Mouse:session intercept to conservatively estimate interactions between these lower-level variables with any higher-level variable (Heisig and Schaeffer, 2019). To test hypotheses on the relationship between bout characteristics (e.g., bout duration, onset time) and neuronal calcium signal, we added scaled and centered characteristics as fixed variables to the signal model for each hypothesis in Experiment 1 (“signal-characteristics models”). Including consumption in the signal model significantly increased fit (Log Likelihood Test, χ^2^_12_ = 35.329, *p* = 0.0004), as did inclusion of bout duration (Log Likelihood Test, χ^2^_12_ = 187.046, *p* < 0.0001) and position (Log Likelihood Test, χ^2^_24_ = 83.031, *p* < 0.0001). For Experiment 2, which had fewer within-subject replications per solution, including Solution as a signal MLMM random slope led to singularity, highly problematic multicollinearity, and model convergence issues. Therefore, we included Time as the only random slope for Experiment 2.

## Results

### Photometry Control Signal Correction

The experimental design of the current studies is described in **Figure 1**. While processing the fiber photometry data, we observed a known problematic artifact where the control 405 nm signal dropped during high activity in the 470 nm GCaMP7f signal (**Figure 2A**), resulting from 470 nm signal “bleed-through” (Simpson et al., 2024) and causing artificial inflation of the normalized calcium signal. We developed a machine-learning B-spline correction (JCBM) to eliminate the influence of this artifact on the normalized GCaMP7f signal. As shown in **Figure 2B**, the JCBM method eliminated the appearance of 405 nm downward deflections. We then tested the JCBM method’s effect on the 470 nm signal during the deflection artifacts and found that the Standard method, as expected, significantly inflated the mean 470 nm signal during deflections (*p* < 0.0001), the Robust method suppressed mean signal (*p* < 0.0001), and the JCBM method neither inflated nor suppressed mean signal (*p* = 0.6794; **Figure 2C**). Full factorial statistical analyses are shown in *Supplement B*. The JCBM method, thus, effectively eliminated the 405 nm artifact’s influence on the normalized 470 nm signal.

**FIGURE 2.**
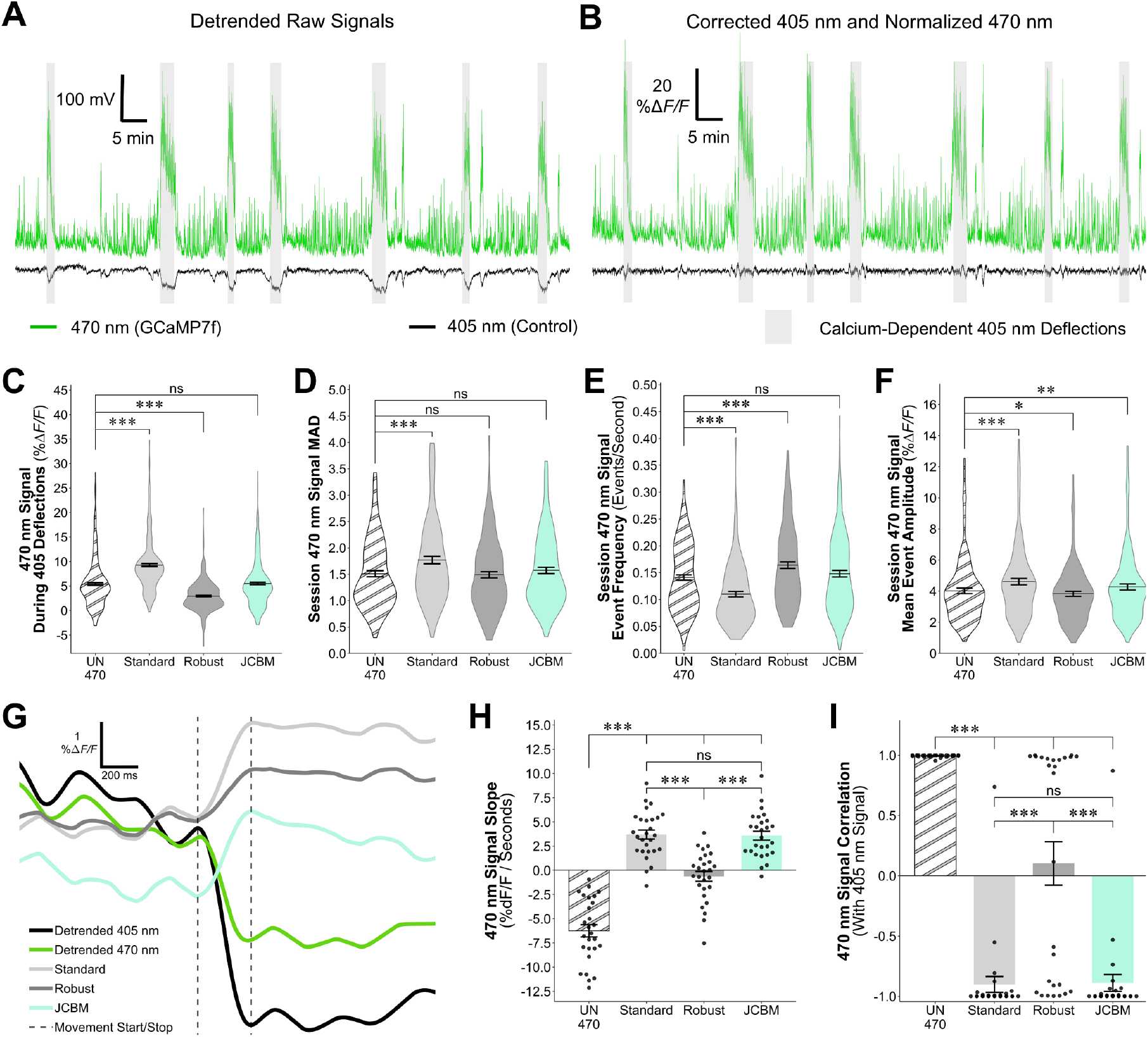
Representative raw signal traces (**A**) prior to signal normalization where 470 nm signal “upstates” were coupled by 405 nm signal downward deflections, indicating calcium-dependency in the intended isosbestic control. (**B**) The same traces after 405 nm signal correction using the Joint method and 470 nm signal normalization. Evaluation of 470 nm signal dynamics showed that normalization using the Standard method inflated (**C**) signal during 405 nm signal deflections, increased (**D**) session variation (MAD), suppressed (**E**) detected peak frequency, and inflated (**F**) detected peak amplitude. Correction of the 405 nm signal using the Robust method had varying but generally opposite effects on 470 nm signal dynamics as the Standard method. Correction using the Joint method showed less signal inflation/suppression and preserved more 470 dynamics than other methods. Evaluation of movement artifact normalization using different methods (**G**, example traces) showed that the Joint method was equal to the Standard method in removing signal distortion (**H**, slope of artifact; **I**, correlation of signals). Graphs display mean ± SE. *ns*, not significantly different; *, *p*< .05; **, *p<* .01; ***, *p*< .001.

Next, we investigated if the JCBM method compromised the normalized signal outside of these deflections. As shown in **Figure 2D**, the Standard method inflated the mean absolute deviation (MAD) of the 470 nm signal (*p* < 0.0001), whereas Robust and JCBM methods did not differ significantly from 470 UN (*ps* = 0.6049 and 0.0845, respectively), indicating that these methods preserve unnormalized 470 nm signal variation. When we investigated normalization method effects on frequency and amplitude of GCaMP7f events (**Figures 2E-2F**), we found that the Standard method decreased frequency and increased amplitude relative to unnormalized signal (*ps* < 0.0001). The Robust method increased frequency and decreased amplitude compared to unnormalized signal (*ps* = 0.0279 and < 0.0001). The JCBM method did not significantly affect event frequency (*p* > 0.05), but increased amplitude (*p* = 0.0037). Notably, event amplitude using the JCBM method was increased significantly less than the Standard method (*p* = 0.0005). These results suggest that compared to the alternative methods tested, the JCBM method produced the least distorted, normalized 470 nm signal.

A source of extraneous fluctuations in the GCaMP7f signal is from movement artifacts produced by shifts in the brain relative to the optic fiber and movements of patch cords and their connectors (Simpson et al., 2024). To test the performance of the JCBM method in removing movement artifacts, we induced robust movement artifacts and measured the slope of the 470 nm signal and its correlation to the 405 nm signal. Our rationale in analyzing these measures was that the slope of the change in signal during an unnormalized movement artifact should be large but closer to zero when adequately normalized. Likewise, we reasoned that the 470 nm signal should be highly correlated with the 405 nm signal during an unnormalized movement artifact and thus adequate normalization should relax the correlation between the two signals. An example signal trace showing an initial drop in signal during a movement artifact is shown in **Figure 2G** with overlaid signal traces from each normalization method. All three methods showed significant reversals in negative 470 nm signal slope (**Figure 2H**; *p* < 0.0001). The Standard and JCBM methods were the only methods whose slopes did not differ from each other (*p* = 0.9919). All normalization methods reduced the positive correlation between the 470 and 405 nm signal during the movement artifact (**Figure 2I**). Standard and JCBM did not differ in this effect from each other (*ps* < 0.0001). Although the 470-405 correlation for the Robust method was near zero, there was high variation in the correlations with half of the correlation near +1 and half near −1. Collectively, this suggests that the JCBM method performs at least as well as the Standard method in correcting movement artifacts. The JCBM method, thus, was used to normalize photometry data from both experiments.

### Experiment 1: CeA^Dyn^ Calcium Transients Around Alcohol Drinking Are Larger Compared to Water and Sucrose Drinking

The experimental data structure is presented in **Figure 3A**. To account for hierarchical data, account for random differences between repeated sessions and bouts within mice, and to provide better statistical power to test relationships between the primary factors of interest, we employed a MLMM approach with random slopes and intercepts (see **Table 1**). A priori, we were interested in the differences in CeA^Dyn^ neuron activity around a bout of drinking based on which solution was being consumed and the biological sex of the mouse. We recorded CeA^Dyn^ calcium activity during voluntary access to alcohol (EtOH), water, or 0.5% sucrose (**Figure 1A**, Experiment 1). Individual bout signals and group averages are shown in **Figure 3B.** The average calcium signal within a 30 second window before and after a drinking bout for EtOH, water, or sucrose is shown in **Figure 3C**. Helmert model contrasts for solution allowed us to test whether sucrose signal was different than water and then whether EtOH was different than the average of both water and sucrose. There was a significant interaction between solution and time where EtOH had a larger change in signal around a drinking bout than sucrose and water drinking bouts jointly (*p* < 0.0001). Post hoc analysis confirmed that the signal change around a bout of EtOH drinking was significantly higher than the signal around drinking water or sucrose (*ps* < 0.0001), indicating that CeA^Dyn^ neurons show a more robust engagement associated with a bout of alcohol drinking than either water or sucrose. No other effects were significant (*p* > 0.1233), including those related to sex. The estimated means, collapsed across sex, and corresponding confidence intervals of this model are shown in **Figure 3D**.

**FIGURE 3.**
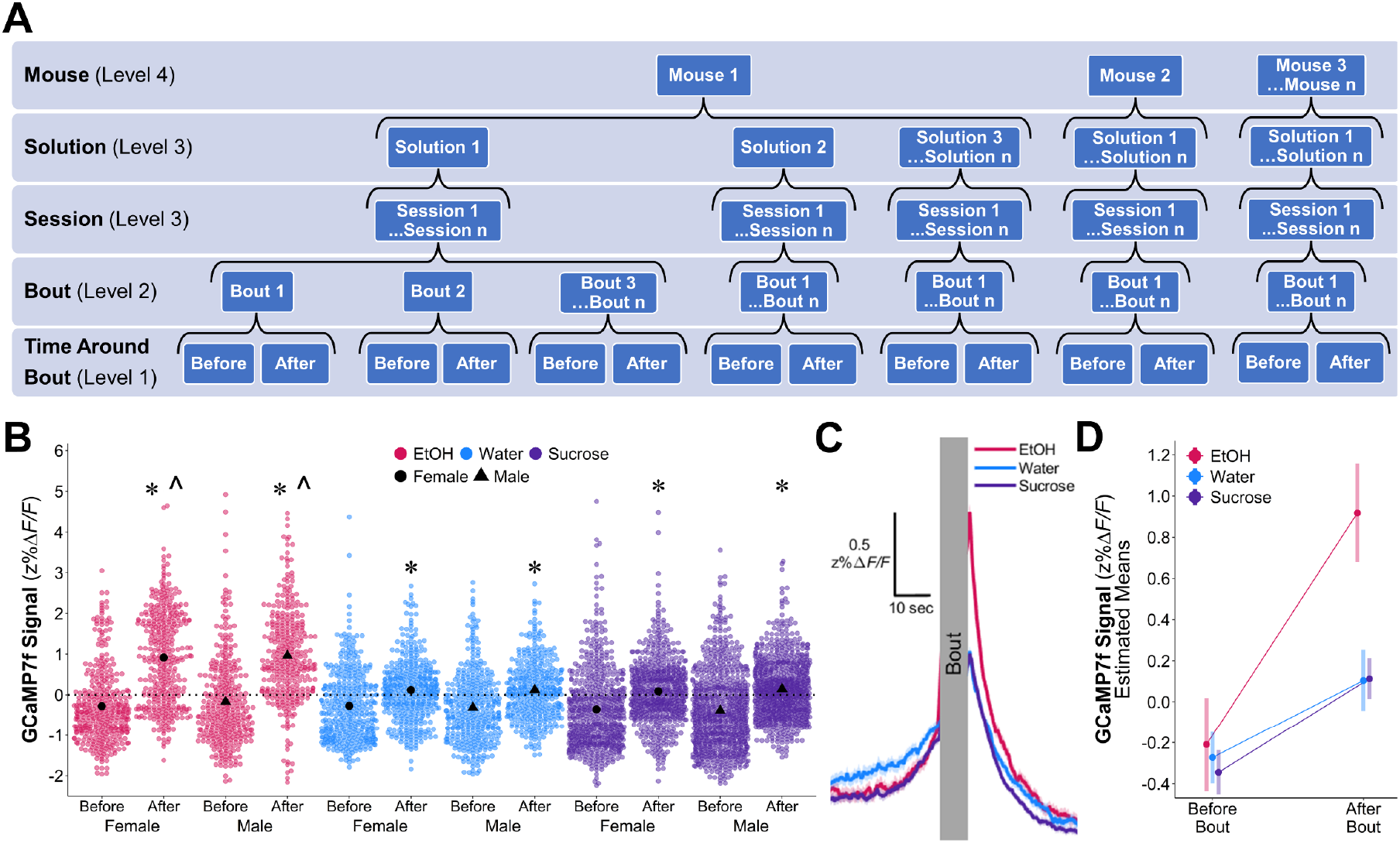
(**A**) Data structure and level assignment for multilevel linear mixed modeling (MLMM). (**B**) GCaMP7f signal (z%Δ*F/F*) 1 sec before and after individual drinking bouts for EtOH (red), water (blue), and 0.5% sucrose (purple) with group means (black circles: females; black triangles: males). (**C**) Mean ± SE GCaMP7f signal (z%Δ*F/F*) traces 30 seconds before and 30 seconds after bouts, collapsed across sex. Signal-outcome MLMM (**D:** estimated means ± 95% CI) showed signals after bouts were significantly higher than before bouts signal for all solutions, but that EtOH showed a more pronounced increase in signal compared to water and sucrose. There were no effects of sex, thus average traces and estimated means are shown collapsed across sex. *, significantly higher than respective before-bout signal, *p* <.05; ^, significantly higher signal increase around bout compared to water and sucrose.

### Drinking Characteristic Differences and Relation to Signal

We next hypothesized that the alcohol- and bout-specific calcium signal in CeA^Dyn^ neurons may be explained by differences in drinking behavior (e.g., bout duration, volume consumed, etc.) for EtOH compared to water and sucrose. To test this hypothesis, we first characterized drinking behavior differences between the solutions with the intent of discovering candidate variables to include as predictors in the signal outcome model. Despite a lack of sex-dependent effects on signal from the previous analysis, sex nevertheless could have mediated relationships between drinking behavior and signal. Thus, we investigated the impact of both sex and solution on drinking characteristics using an outcome model for each drinking characteristic. After fully describing the characteristic with its own model, we then included the characteristic as a fixed factor in the signal-outcome model and investigated whether the characteristic was significantly associated with GCaMP7f activity. We focused on these characteristics: consumption (mL/kg) at the session level (Level 3) and bout duration and onset time (position) at the bout level (Level 2).

#### A. Session Consumption

The volume of each solution consumed (mL/kg) is shown in **Figure 4A**. The estimated means for the consumption model are shown in **Figure 4B**, demonstrating that there was an interaction between sex and solution. Females drank less EtOH relative to sucrose (*p* < 0.0001) and water (*p* = 0.0003), yet males drank less EtOH than sucrose (*p* = 0.0003) but not water (*p* = 0.6920). Each sex consumed similar amounts of water and sucrose (*ps* > 0.1338); however, female mice drank more water and sucrose than males (*ps* = 0.0267, 0.0461). For EtOH specifically, there was no consumption difference between males and females (*p* = 0.9622).

**FIGURE 4.**
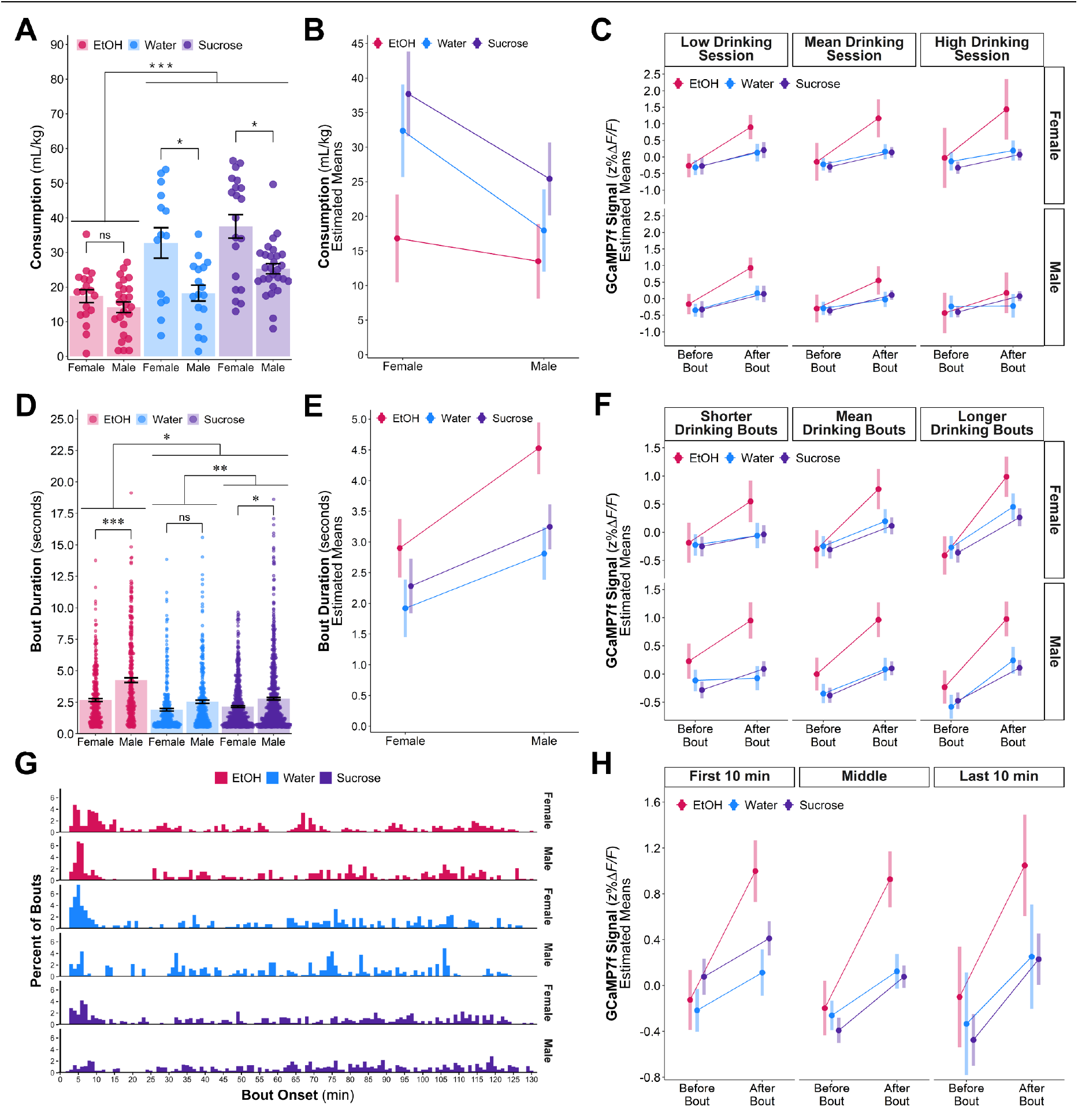
(**A**) Mean ± SE consumption of each solution (mL/kg), divided by sex. Consumption MLMM (**B**: estimated means ± 95% CI) showed that females generally drank more than males, except with EtOH where both sexes drank similar amounts. Both sexes also drank more water than EtOH and more sucrose than both EtOH or water. Signal-Consumption MLMM (**C**: estimated means ± 95% CI for females (top), males (bottom)) showed that during sessions with low EtOH or water consumption (−1 SD), males, but not females, show an increase in signal around a drinking bout. Neither sex showed a consumption-associated change in signal for sucrose drinking bouts. (**D**) Mean ± SE bout duration (sec) by solution and sex. Duration MLMM (**E**: estimated means ± 95% CI) showed that both sexes drank EtOH in longer bouts than water and sucrose; males more so than females. Signal-Duration MLMM (**F**: estimated means ± 95% CI) showed that longer bout duration (+1 SD) was associated with a greater signal change around a bout, however, even at equally high durations, EtOH-associated signal remained higher than water/sucrose. (**G**) Histogram of bout onset for each solution and sex. Signal-Position (onset) MLMM (**H**: estimated means ± 95% CI) showed that sucrose signal decreased on average across the 2-hr session even while the increase around a drinking bout was amplified. In contrast, the signal increase around a bout did not change for water nor EtOH within sessions. *ns*, not significantly different; *, *p* <.05; **, *p* <.01; ***, *p* <.001.

Since the unique signal change in CeA^Dyn^ neurons around an EtOH drinking bout might be explained by EtOH’s lower volume consumption, we investigated whether lower consumption led to a larger increase in signal around a bout compared to high consumption. Individual mouse mean-centered consumption was included as a predictor in the signal-outcome model to compare signals between low and high volume drinking sessions. Adding consumption to the model introduced new interactions with sex (**Figure 4C**). There were interactions between consumption and time around a bout (*p* = 0.0103), consumption and sex (*p* = 0.0397), and a three-way interaction between consumption, time, and sex (*p* = 0.0019). There was also a 4-way interaction between consumption, time, sex, and solution (*p* = 0.0240). Post hoc analyses of these interactions revealed that the signal change around a bout was significantly higher in low consumption sessions for EtOH and water in males (*ps* = 0.0005, 0.0021) while there were no differences for these solutions in females (*ps =* 0.0840, 0.3688). These results partially support the hypothesis that, at least in males, lower consumption of EtOH is associated with higher changes in CeA^Dyn^ neuron activity. Consumption did not, however, appreciably change signal increase around a bout for sucrose in either sex (*ps* > 0.5571), indicating that if this hypothesis is true, it is both sex- and solution-specific. These results indicate that the amount of consumption does not easily explain greater activity of CeA^Dyn^ neurons around EtOH drinking.

#### B. Bout Duration

At the bout level, bout duration (**Figure 4D**) showed an effect of solution where sucrose bouts were longer than water bouts (*p* = 0.0011) in both sexes due to absence of an interaction (*p* = 0.7630). EtOH bouts were longer than water and sucrose jointly (*p* < 0.0001) but there was also a significant interaction of this comparison with sex, where males had a greater relative increase in EtOH bout durations (versus water/sucrose) than females (*p* = 0.0027). Within EtOH and sucrose but not water, males drank in longer bouts than females (*ps* < 0.0001, 0.0175, 0.0669). The estimated means for the bout duration model are shown in **Figure 4E**. Altogether, these results suggested that under these conditions mice drink EtOH in uniquely longer bouts than water and sucrose and that males and females drink EtOH in different length bouts, despite consuming similar EtOH amounts at the session level.

We then investigated the relationship between signal and bout duration, hypothesizing that if longer bout durations explained the unique CeA^Dyn^ response to EtOH drinking, a longer bout duration would be associated with higher bout-related signal. Bout duration interacted with the effects of time and sex individually such that signal change around a bout was higher with longer drinking durations (*p* < 0.0001) and when males drank longer than usual, on average their increase in signal around a bout was smaller relative to that of females (*p* < 0.0001). Further examination of the estimated means for each sex as a function of bout duration and solution (**Figures 4F**) showed that the change in signal around a bout increased for all solutions in both sexes as an animal drinks longer than they usually do (*ps* < 0.0001, 0.0053 male sucrose), but that there is no interaction between bout duration with the relative differences in bout-related signal between EtOH and water/sucrose (*p* = 0.3302). At long bout durations, EtOH-associated signal around a bout is still greater than that of other solutions (*ps* < 0.0001, except male EtOH-water *p* = 0.0054). While bout duration has a positive relationship with signal, this characteristic cannot explain EtOH’s unique engagement of CeA^Dyn^ neurons.

#### C. Bout Onset (Position)

When we examined a histogram of bout onset across the session for each solution (**Figure 4G**), we found that for each group (except for males drinking sucrose), the highest proportion of licking bouts occurred within the first 5-10 min. Mice are known to “frontload” EtOH, and this is thought to relate to EtOH vulnerability (Ardinger et al., 2022). If early drinking of EtOH was related to the unique change in CeA^Dyn^ signal, this would have important implications for the development of maladaptive EtOH drinking. We were interested, thus, in whether the temporal occurrence (“position”) of the bout within the 2-hr session was predictive of signal change magnitude around a bout and if the signal changed over time, perhaps as a response to satiation or intoxication. We added position within the session to the signal-outcome model. We hypothesized that EtOH would show a prominently higher signal increase around a bout within the first 10 min and this unique signature would habituate over the 2-hr drinking sessions. After including position in the signal outcome model, the estimated mean signals for bouts within the first/last 10 min and the time in between are shown in **Figure 4H**. There was significant effect of position alone where average signals within the first 10 min were higher than the rest of the session (*p* = 0.0184), irrespective of solution and time around bout. With position in the model, there was now an effect of sex such that males had slightly higher average signals than females (*p* = 0.0454), but there were no interactions of any factor with sex. Using post hoc comparison of the first and last 10 min within solution, we found that the sucrose signal change around a bout increased significantly across the session (*p =* 0.0152). Since there was no interaction of position with the change in EtOH signal around a bout relative to sucrose/water (*p* = 0.2758), the relatively larger EtOH signal change neither habituated nor amplified across the 2-hr session. We also confirmed from post hoc comparison that there was no significant difference in the signal change around a drinking bout between the first and last 10 min for neither EtOH (*p* = 0.9222) nor water (*p* = 0.3882). As with total session consumption and bout duration, temporal occurrence of the bout also did not explain the unique signal for EtOH.

### Experiment 2: CeA^Dyn^ Calcium Transients in Response to Negative Valence, Valuation State, and Taste Do not Recapitulate Unique Alcohol-Associated Response

To explore other potential explanations for why alcohol drinking would be related to a larger signal increase, we recorded CeA^Dyn^ calcium activity during voluntary access to a battery of different solutions (**Figure 1A**, Experiment 2) intended to test different hypotheses about what this unique EtOH-associated signal encodes.

#### A. Quinine – Bitterness/Aversiveness

One hypothesis for why alcohol would be associated with a unique signal in CeA^Dyn^ neurons is that alcohol likely has more aversive and/or bitter taste characteristics compared to water and sucrose. To test this possible explanation, we examined CeA^Dyn^ calcium activity during voluntary consumption of varying concentrations of quinine, a known bitter tastant. We gradually reduced the concentration of quinine daily until it reached 90 µM, as this was where most mice of both sexes began to consume measurable amounts. We observed that before lowering the concentration to 90 µM quinine, females begun consuming quinine at higher concentrations than males (1 mM vs 125 µM, not shown) suggesting that females may be less behaviorally sensitive to quinine. Consumption (mL/kg) during 90 µM “low” quinine access (**Figure 5A-5B**), contrary to expectation, did not differ significantly from water (**Figures 5A-5B**; *p* = 0.4414). **Figures 5C-D** shows CeA^Dyn^ GCaMP7f signal for individual bouts and 30 sec before and after a bout, and the estimated means and CI are shown in **Figure 5E**. Analysis showed that quinine did not differ from water in the signal change around a bout (*p* = 0.2023) but was associated with lower average signal than water (*p* < 0.0001). These results suggest that quinine drinking is associated with a general decrease in CeA^Dyn^ activity relative to water. Quinine drinking also does not recapitulate the larger increase in CeA^Dyn^ activity seen around an alcohol drinking bout, suggesting this increase is not due to alcohol aversiveness/bitterness.

**FIGURE 5.**
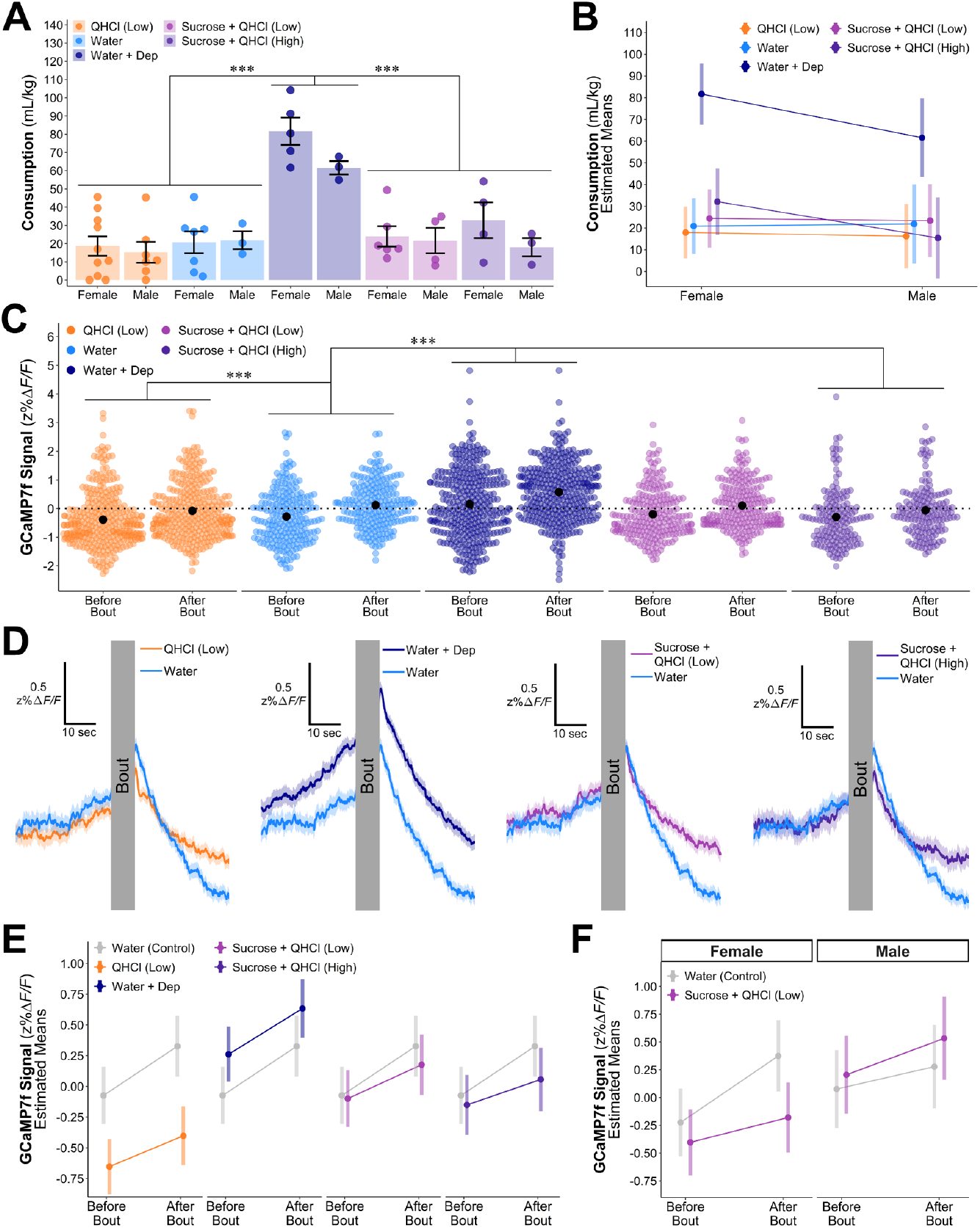
Experiment 2 results for test solutions QHCl (Low), water, water under water deprivation, sucrose + low QHCl, and sucrose + high QHCl. (**A**) Mean ± SE consumption (mL/kg). Consumption MLMM (**B**, estimated means ± 95% CI) showed that male and female mice drank more water under water deprivation than other solutions. (**C**) Individual bout and group mean ± SE signal (z%*ΔF/F*) 1 sec before and after bouts. (**D**) Average signal 30 seconds before and after bouts, divided by solution alongside water (light blue). Signal MLMM (**E**, estimated means ± 95% CI) showed that, irrespective of time around bout, QHCl (Low) and sucrose + high QHCl drinking were associated with lower average signal while drinking water under water deprivation was associated with higher average signal. No test solution recapitulated the unique EtOH signal around a drinking bout from Experiment 1. Sucrose + low QHCl (**F,** estimated means ± 95% CI) showed sex-dependent differences in signal change around a bout. ns, not significantly different; *, *p* <.05; **, *p* <.01; ***, *p* <.001.

#### B. Water Under Deprivation – Altered Value

To determine how a change in motivational value of a solution alters CeA^Dyn^ activity, we recorded CeA^Dyn^ calcium transients during water drinking sessions with and without water deprivation. As expected, mice drank significantly more water under a water-deprived state than water in a non-deprived state (*p* < 0.0001), confirming an increased motivational value of water by deprivation (**Figures 5A-5B**). Analysis indicated that water drinking in a deprived state did not produce significantly different signal change around a drinking bout (*p* = 0.7935; **Figures 5C-E**) but was associated with significantly increased average signal (*p* < 0.0001). These results suggest that increasing value is associated with increased general CeA^Dyn^ activity but not increased signal change associated with a drinking bout.

#### C. Sucrose with Quinine – Sweet and Bitter Combination

Alcohol has predominantly bitter taste characteristics for humans and rodent models alike at lower concentrations, but at higher concentrations, can have an even stronger sweet-taste component. Thus, mixtures of sweet and bitter are proposed to better mimic the taste of alcohol (see (Mukherjee et al., 2023)). Here, we tested whether a mixture of sucrose and quinine, 0.5% sucrose with either 125 µM “high” or 90 µM “low” quinine, recapitulates an alcohol-like CeA^Dyn^ calcium response due to its similar bitter-sweet characteristics. Comparison of consumption between sucrose + low quinine and sucrose + high quinine (**Figures 5A-5B**) indicated that there was not a difference in consumption between the two mixtures (*p* = 1) nor either mixture compared to water (*ps* = 0.6861 (low), 0.7239 (high)). Signal analysis indicated that there was an interaction of sucrose + low quinine with sex and time around a bout where the difference in signal change around a bout between sucrose + low quinine and water was significantly higher in males than females (*p* = 0.0285; **Figure 5C-5D**, collapsed by sex, **Figure 5F** divided by sex). Interrogation of this interaction showed that the sucrose + low quinine signal change was decreased compared to water for females but not for males (*ps* =0.0308 female, 0.3963 male). These results suggest that bitter-sweet mixtures are associated with changes of CeA^Dyn^ neuron activity around about, but only in females and in the opposite direction from the unique EtOH-associated response of these neurons. Conversely, sucrose + high quinine signal change around a bout was not different than that of water (*p* = 0.1268), nor was there an interaction with sex (*p* = 0.7995). Average sucrose + high quinine signal was significantly lower than water (*p =* 0.0129) without an interaction with sex (*p* = 0.1167), indicating that in both sexes, while the change in signal around a sucrose + high quinine bout is like water, average signal is lower than water. Collectively, these results suggest that the unique EtOH-associated response of CeA^Dyn^ neurons cannot be attributed to bitter-sweet taste.

## Discussion

Dynorphin-expressing neurons in the central amygdala (CeA^Dyn^) have been heavily implicated in mediating excessive alcohol intake and excessive alcohol drinking produced by the interaction of stress and alcohol dependence. The current fiber photometry study demonstrated that EtOH drinking uniquely engages the CeA^Dyn^ neuronal population – there is a larger increase in calcium activity in CeA^Dyn^ neurons associated with a bout of drinking alcohol than drinking water and sucrose. Our analyses tested whether drinking characteristics that differ for alcohol relative to water and sucrose are related to this unique alcohol-associated signal. While these drinking characteristics were associated with modulated signal, none satisfactorily explained alcohol’s association with greater engagement of CeA^Dyn^ neurons. To more deeply test whether the taste and value characteristics of alcohol can recapitulate alcohol’s uniquely increased activity in CeA^Dyn^ neurons, we measured calcium transients around drinking bouts for bitter, bitter-sweet, and water under water deprived conditions and found that none of these conditions convincingly reproduced higher activity specifically around a drinking bout seen with alcohol.

Recently, others have also investigated CeA^Dyn^ neuron activity around alcohol consumption and found that water, sucrose, and alcohol drinking bouts were associated with equally evoked CeA^Dyn^ neuronal activity (Roland et al., 2024). While at first glance these results seem at odds with our findings, there are methodological differences that could account for the differing results. Firstly, in the current studies mice were allowed 2 hr for drinking instead of intermittent 5 second periods. Shorter momentary access to solutions could change the urgency/motivation to respond and therefore modify CeA^Dyn^ neuron activity. While most bout durations were below 5 seconds for each solution in our study, there were a significant number of much longer bouts (~10-15 seconds), particularly with alcohol, when mice were allowed to voluntarily choose when, how long, and how often to drink. These bout sampling differences combined with different activity measurements during drinking could have contributed to differing results. Additionally, during each of our 2-hr sessions, mice were only allowed access to a single testing solution, whereas in Roland et al., mice received access to multiple solutions in a randomized sequence. It is unclear what impact the novelty or salience of rapidly switching solutions would have on relative evoked signals, but current results suggest that novelty does not have an important influence on measured CeA^Dyn^ activity. The authors also tested a 10-fold higher concentration of sucrose (5% vs 0.5%) which could have reduced signal differences between sucrose and alcohol. Indeed, taste of sucrose concentrations around 5% may better generalize to alcohol solutions based on previous studies (Blizard, 2006). The different concentrations of alcohol used between our studies (15% vs 20%) may also have impacted these relative signal differences and sweet/bitter taste characteristics. Another major difference between studies is that Roland et al. water deprived mice to encourage drinking during short test periods, whereas our mice were not water deprived. While the authors mention they tested mice in water non-deprived states and saw no differences in mean responses with water or alcohol in the global CeA, it is unclear whether water deprivation in this context would specifically influence signal in the CeA^Dyn^ subpopulation, which showed several notable differences from the pan CeA recordings in their studies. The current studies provide evidence that water deprivation increases the activity of CeA^Dyn^ neurons generally during water drinking and thus it is possible that deprivation state-induced activity shifts or deprivation-induced stress might obscure differences between solutions. In Roland et al., the CeA^Dyn^ subpopulation response did not habituate during test sessions, in contrast to pan CeA responses. While the current studies did not measure pan CeA responses, our results also showed no habituation of the CeA^Dyn^ subpopulation response to alcohol within session. On a broader time scale, Roland et al. showed that CeA^Dyn^ alcohol-associated responses were still present after 3 weeks of continuous alcohol access, in agreement with the current studies. Finally, Roland et al. CeA^Dyn^ responses showed a greater quinine associated responses in females versus males, a sex difference not found in the current study. Despite these differences, both studies’ results agree that CeA^Dyn^ activity increases around a bout of alcohol drinking and does not habituate over time.

Our finding that alcohol is associated with larger increases in CeA^Dyn^ neuron activity than other solutions has important implications for the role of these neurons in alcohol drinking and on alcohol effects on behavior governed by the CeA^Dyn^ population. A possible explanation of alcohol drinking’s unique association with CeA^Dyn^ activity is that this activity encodes innate alcohol-specific reinforcement, which could lead to excessive alcohol intake with repeated exposure. While this explanation might be attractive because it suggests alcohol’s reinforcement is “hard-wired” or innate, there is a great discussion in Janiak et al. (Janiak et al., 2020) that suggests that higher alcohol metabolism evolves in animals that have a greater fruit and nectar component of their diet to help counter intoxication from consumption of fermented fruits. In this light, the well-documented high alcohol metabolism of *Mus musculus* suggests that this species evolved to deal with alcohol and counter intoxication because of a high fruit diet. Thus, any reinforcement signal evoked by alcohol may better reflect the co-occurrence of alcohol with highly palatable fruits. This is of course purely speculative and would need to be empirically tested. More likely, we assert that there is a yet untested physiological action of alcohol that evokes increased activity in CeA^Dyn^ neurons. We observed that increased calcium transient magnitude was present in the first drinking bout for mice that were consuming alcohol for the first time (see *Supplement C*). Alcohol, thus, innately engages these neurons in a unique manner even prior to experiencing its intoxicating effects and strengthens the possibility that there is a purely sensory explanation for the distinct response to alcohol. When we tested solutions thought to share taste characteristics with alcohol, however, drinking bouts of these solutions did not show greater CeA^Dyn^ neuron signal increases around a bout than water. Oral chemesthetic solutions were not tested in the present study so this characteristic remains a possible sensory explanation for the current results. Alcohol, but not water nor sucrose, causes chemesthetic sensations like burning, tingling, and drying, which in humans is negatively associated with frequency of alcohol drinking (Nolden and Hayes, 2015). These chemesthetic sensations are thought to help protect animals from environmental irritants or toxins in the case olfaction does not provide sufficient cues (see (Green, 2012)). Future studies can test the contribution of chemesthesis to alcohol-associated CeA^Dyn^ neuron possibly by measuring in vivo activity of these neurons while manipulating the concentration of alcohol to change the intensity of these chemesthetic sensations or test other solutions with chemesthetic characteristics (i.e. menthol).

Another possibility is that the unique alcohol-evoked activity in CeA^Dyn^ neurons reflects a response to the distinct taste characteristics of alcohol relative to water and sucrose, but that the current study did not test relevant sucrose-quinine mixtures. Alcohol is reported to exhibit both bitter and sweet tastes in humans, with bitter taste dominating at lower alcohol concentrations (≤16%; (Nolden and Hayes, 2015)). Though we cannot assess the subjective experience of taste in mice directly, one study cleverly used stimulus generalization after conditioned taste aversion to examine which taste is perceptually most like alcohol (Blizard, 2006). They reported that both quinine and sucrose taste aversion generalized to alcohol and vice versa. While we do not see a similar increase in signal around a bout of drinking sucrose, quinine, and sucrose-quinine mixtures as alcohol, we only tested one concentration of sucrose and two similar concentrations of quinine, which together may not collectively be perceptually comparable to 20% alcohol. The conditioned taste aversion study described used much more concentrated sucrose (200 mM or 6.8%) and quinine (1 mM) solutions than the current study (0.5% sucrose, 90-125 μM quinine), although the proportion of sucrose to quinine was similar. To fully investigate whether the bitter-sweet taste component of alcohol is associated with unique CeA^Dyn^ activity, future studies should measure CeA^Dyn^ activity during consumption of varying concentrations of sucrose and quinine.

An important caveat to the current results from testing different tastants is that solutions were tested within-subject and thus order and contrast from previous solutions should also be considered. For example, in the second experiment there was a statistically significant difference in the effect of time around a bout between males and females for water, which was used as a control condition (*p =* 0.0378). There was not a sex difference in CeA^Dyn^ activity around a water drinking bout in the first experimental context (*p* = 0.8275), indicating that the second experimental context sex-specifically influenced water-associated CeA^Dyn^ activity. As the sexes showed indications of different behavioral responses to quinine, the previously presented solution to water, it is possible that this history sex-dependently affected the activity associated with the transition to water drinking. We included sex as a variable and investigated all interactions with sex to adequately control for a potential difference in the control conditions, but the possibility remains nevertheless that these influenced some of the sex-specific effects described. Future studies will want to consider potential order effects, especially as they relate to a control condition.

While we saw no indication of habituation of CeA^Dyn^ neuron alcohol-associated responses within three weeks of exposure, we predict that neural activity adaptations would occur with more chronic activation of CeA^Dyn^ neurons by alcohol. Previous work shows that repeated alcohol exposure increases excitability of CeA^Dyn^ neurons, though this effect was sex-specific (males only) and derived from ex vivo electrophysiology. Roland et al. (2024) also investigated CeA^Dyn^ neuronal activity after chronic alcohol using the continuous access paradigm and found no chronic alcohol-induced changes in CeA^Dyn^ neuronal responses (Roland et al. 2024). Future studies can test the impact of long-term alcohol drinking and alcohol dependence on in vivo neuronal activity to better understand CeA^Dyn^ neuron adaptations in both sexes that contribute to AUD-related behavioral phenotypes.

Finally, our current results provide a new normalization method for fiber photometry data to resolve the known issue of channel bleed-through from a control wavelength. These results demonstrated that the new JCBM method sufficiently reduces bleed-through effects, corrects for movement artifacts at least as well as previous methods, and causes less distortion than previous methods in that it preserves the raw signal dynamics of GCaMP7f. Future studies will be able to apply this method to overcome the challenges of channel bleed-through in GCaMP7f fiber and perhaps other photometry approaches.

In summary, alcohol drinking resulted in a larger increase in CeA^Dyn^ activity compared to other solutions (i.e. water and sucrose) in both male and female mice. While drinking amount, taste, and value had a moderating influence on the magnitude of CeA^Dyn^ neuron activity, no condition reproduced the relatively larger alcohol-associated increase in CeA^Dyn^ activity around drinking. The unique engagement of the CeA^Dyn^ neuron population by alcohol could have important implications for alcohol influences on stress and appetitive behavior governed by the CeA. Thus, it is necessary to further study CeA^Dyn^ neuronal activity related to active drinking since engagement of this crucial cell population could explain behavioral adaptations observed in models of alcohol misuse.

## Conflict of Interest

The authors declare no competing financial interests.

## Author Contributions

**Christina L. Lebonville**: Conceptualization, Methodology, Software, Validation, Formal analysis, Investigation, Writing (original and revision), Data curation, Project administration. **Jennifer A. Rinker:** Conceptualization, Methodology, Writing (revision). **Krysten O’Hara**: Conceptualization, Methodology, Software, Writing (revision). **Christopher S. McMahan**: Writing (original and revision), Methodology, Formal analysis. **Michaela Hoffman**: Software, Writing (revision). **Howard C. Becker**: Conceptualization, Writing (revision), Resources, Funding acquisition. **Patrick J. Mulholland**: Conceptualization, Methodology, Software, Formal analysis, Investigation, Writing (revision), Resources, Supervision, Funding acquisition.

**Funding & Acknowledgements**

These studies were financially supported by the Charleston Alcohol Research Center, Charleston, SC, USA (P50 AA010761), the MUSC SCORE Career Enhancement Core, Charleston, SC, USA (U54 DA016511, CLL), the INIAstress Consortium (U01 AA014095, U01 AA020930), VA Medical Research, USA (BX000813, HCB), and National Institutes of Health grants F32 AA029026 (CLL), S10 OD021532 (PJM), R01 AA023288 (PJM), R01 AA026536 (HCB), and K01 AA025110 (JAR). Funders had no input into the design, collection, analysis and interpretation of data, writing of the report and decision to submit the article for publication.

## Supplement A – RStudio Session Information

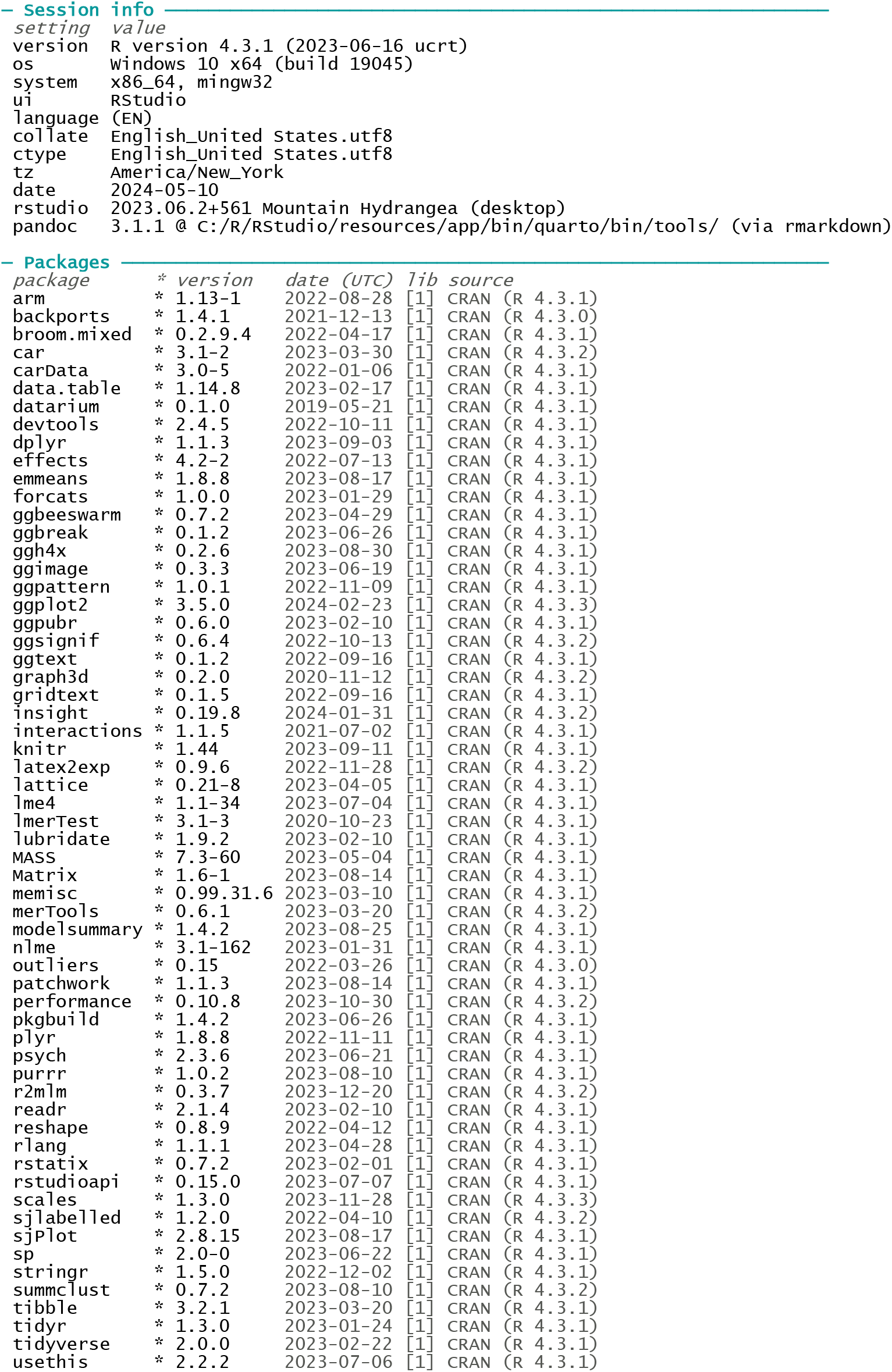
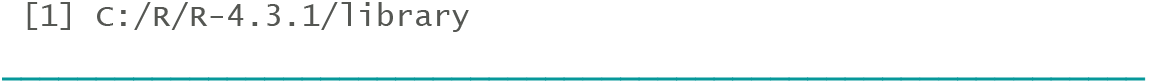

## Supplement B - Full Statistics Report

Abbreviations: E = Effect contrast coded; H = Helmert contrast coded; S = Sucrose; W = Water; Dep = Deprivation; QHCl = Quinine

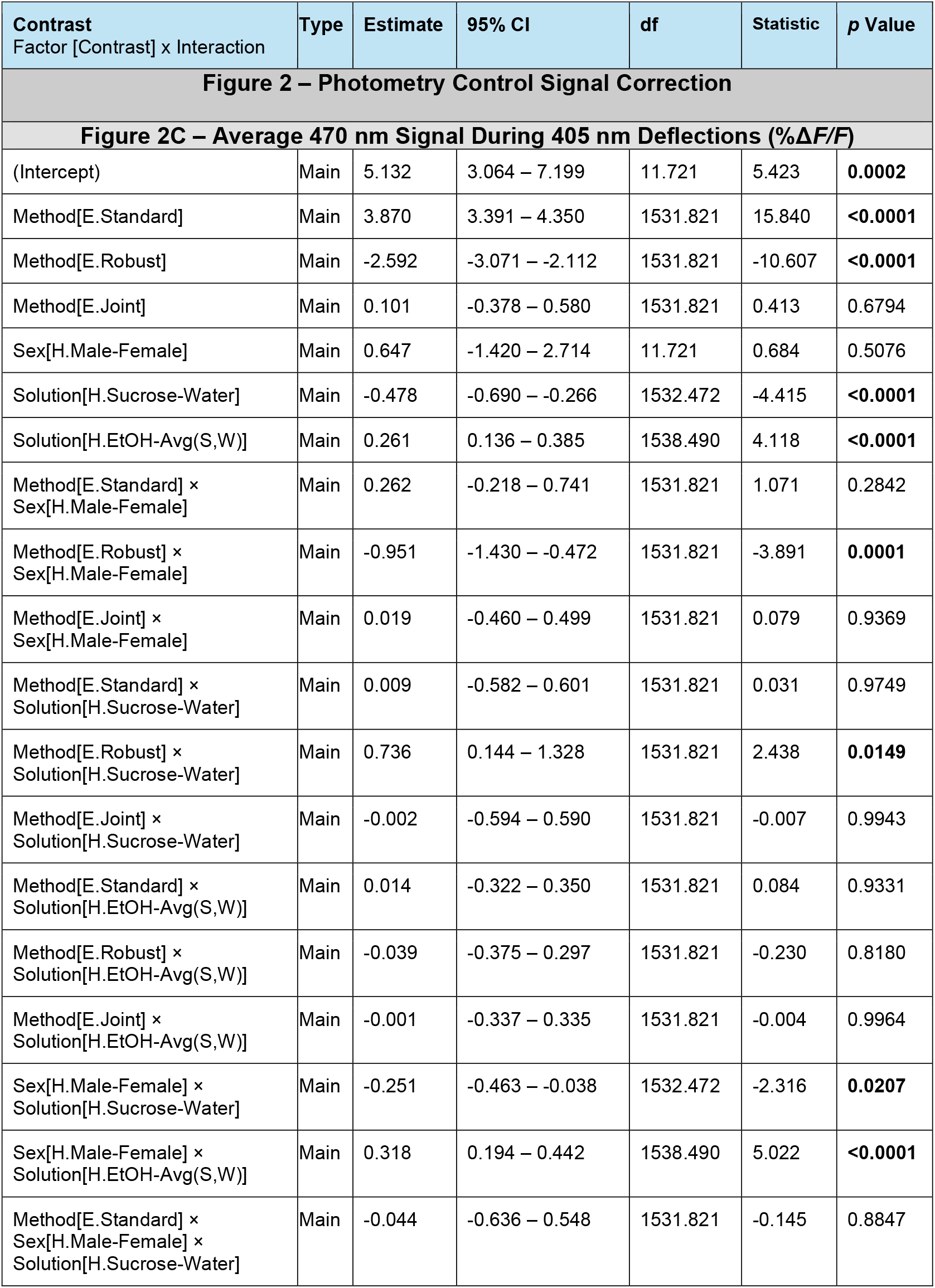
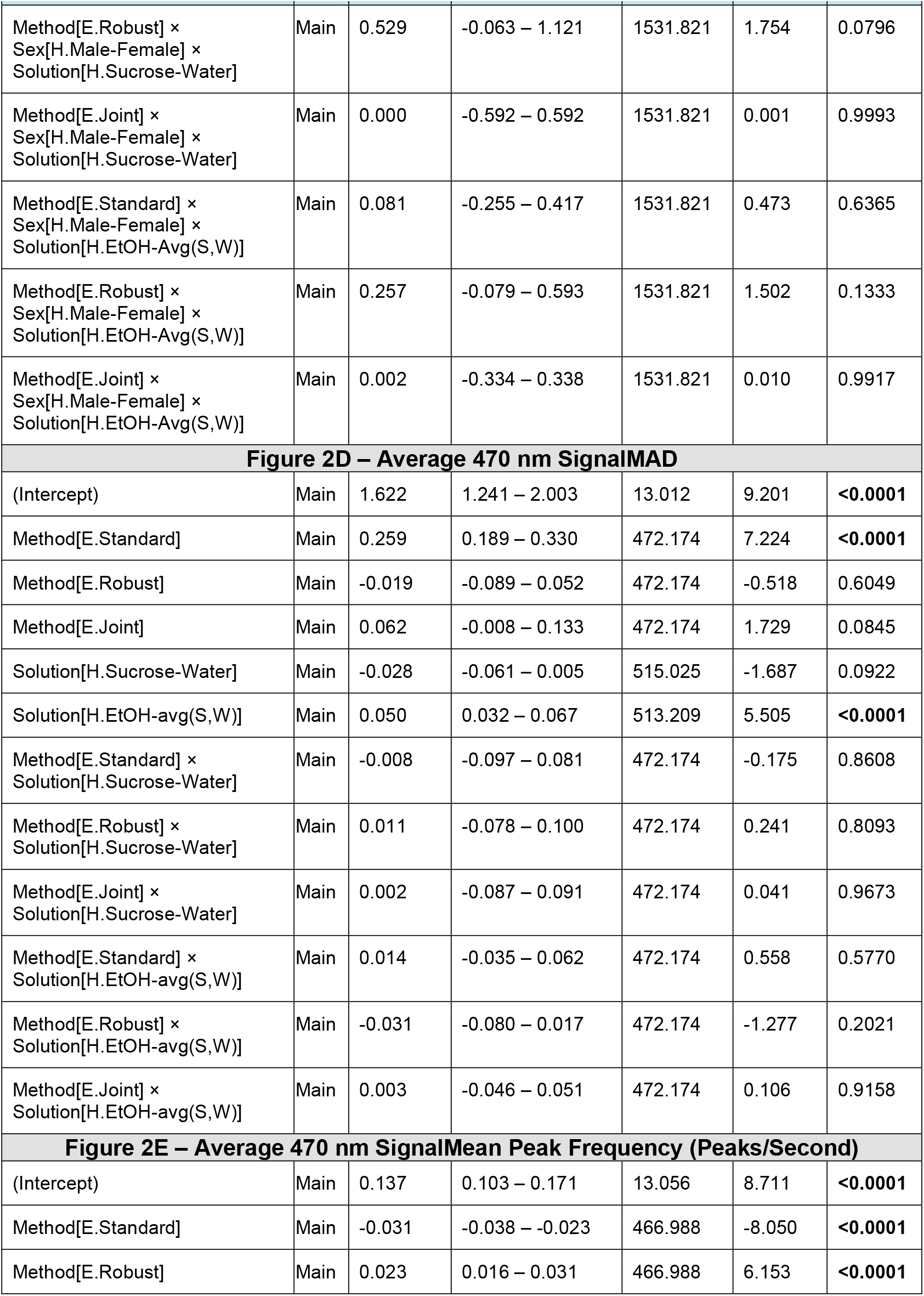
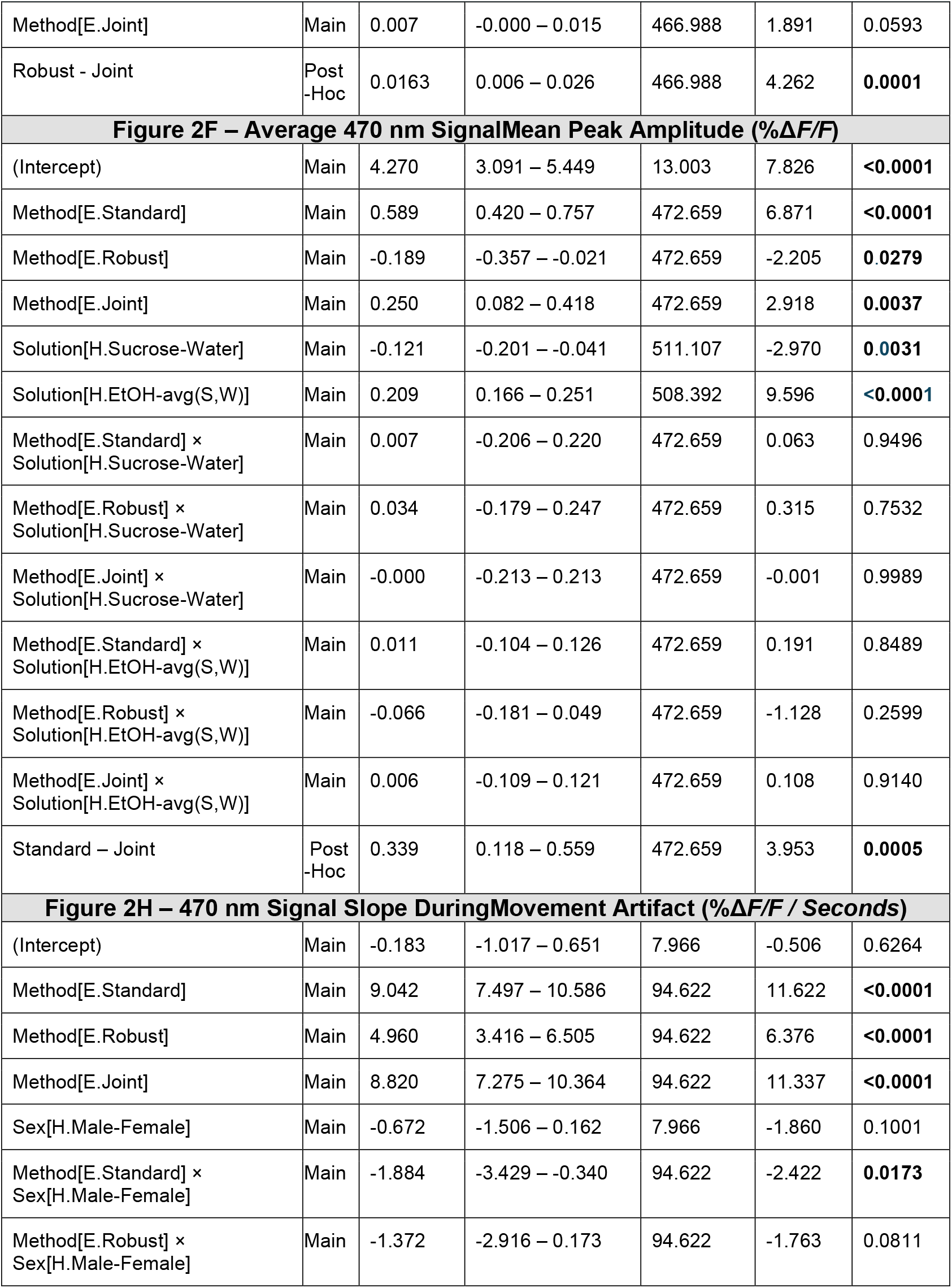
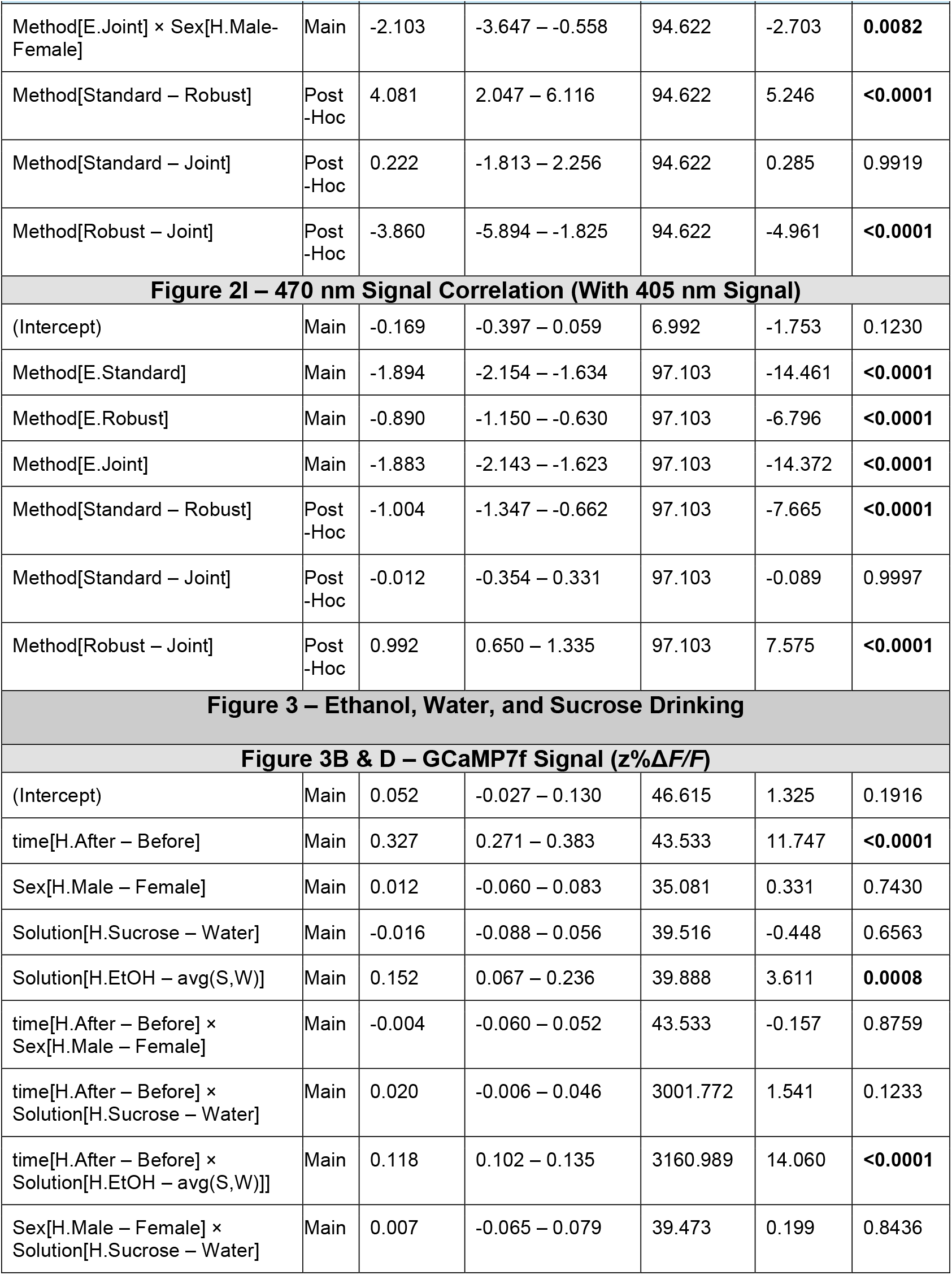
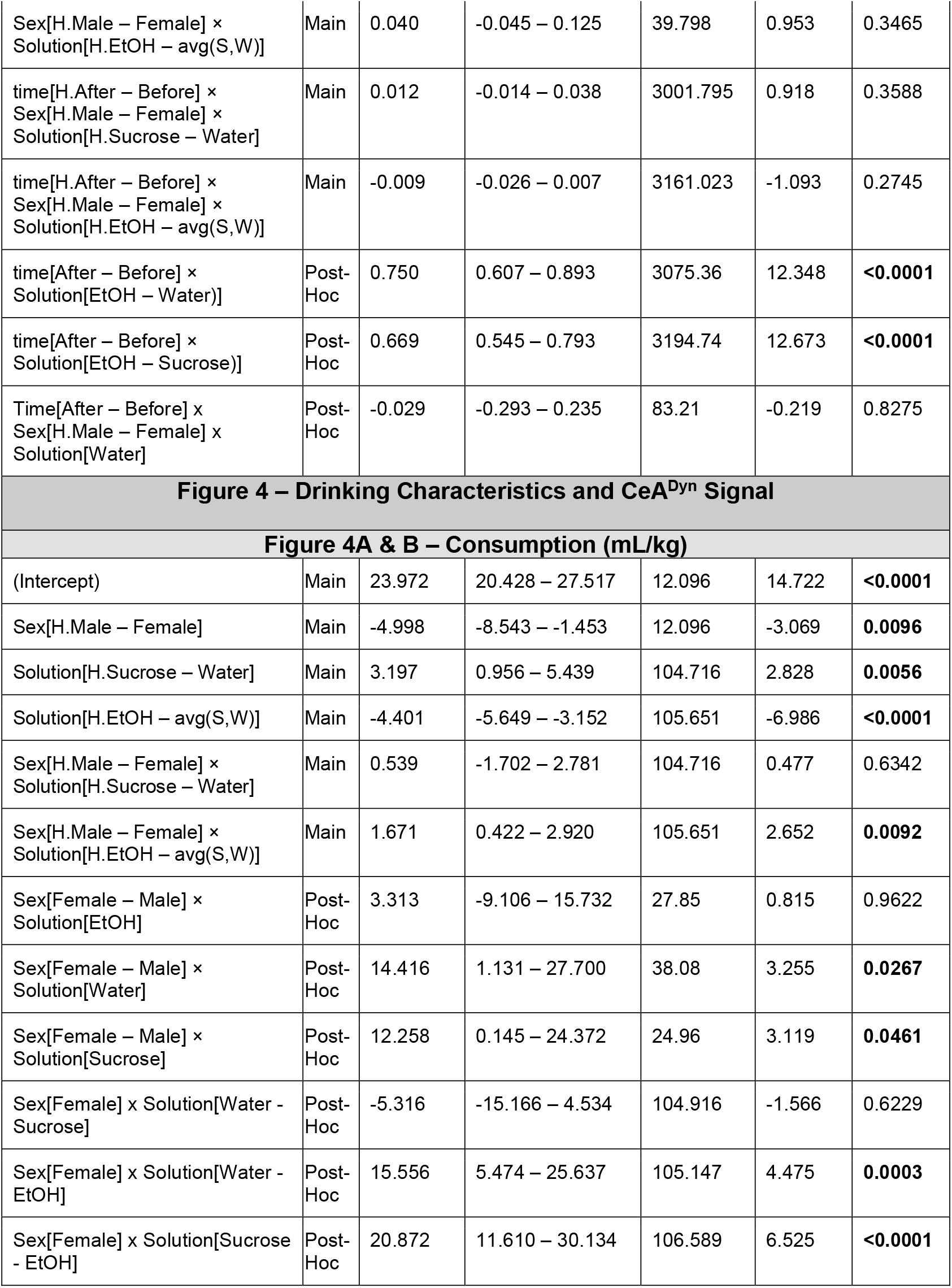
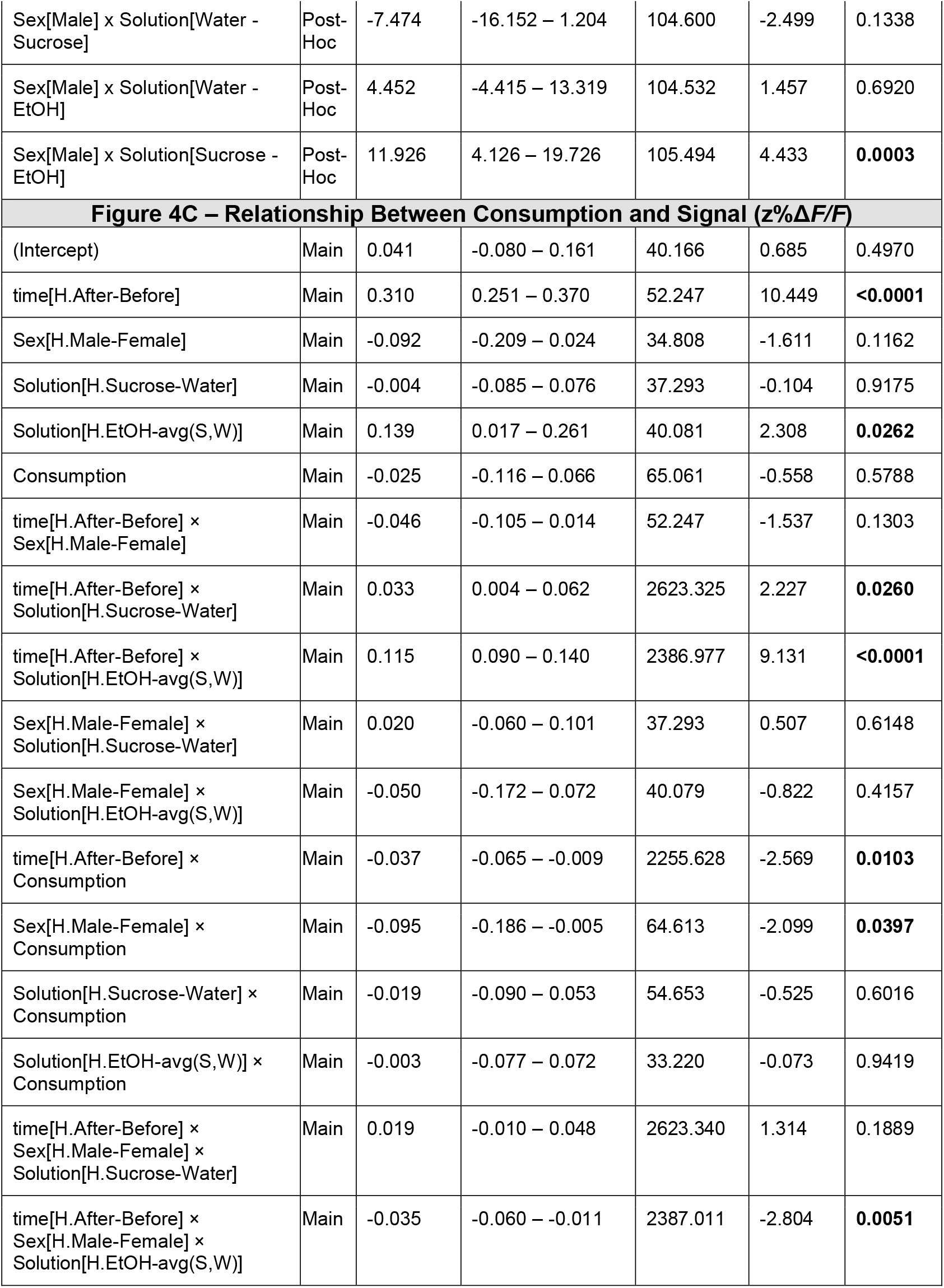
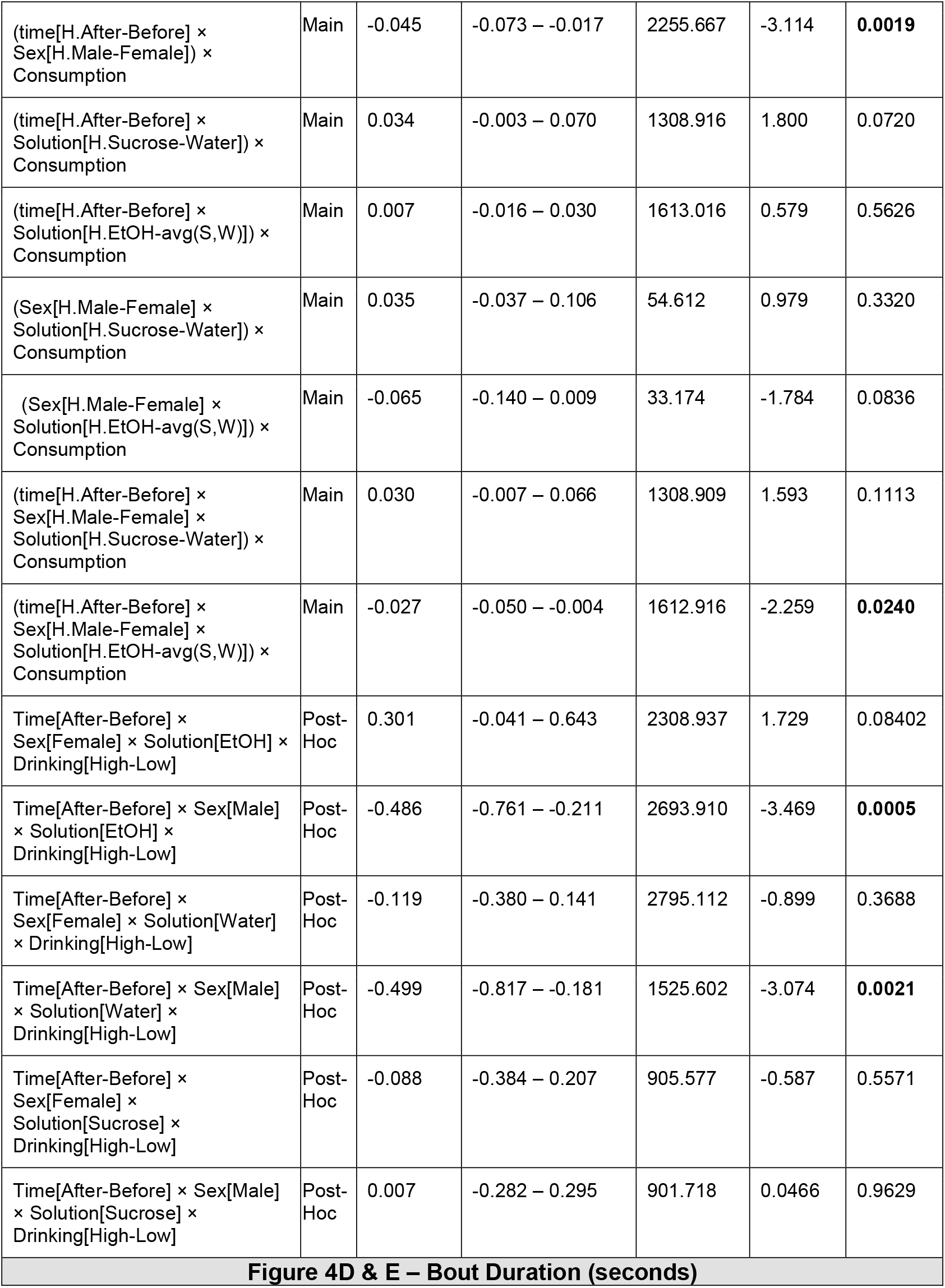
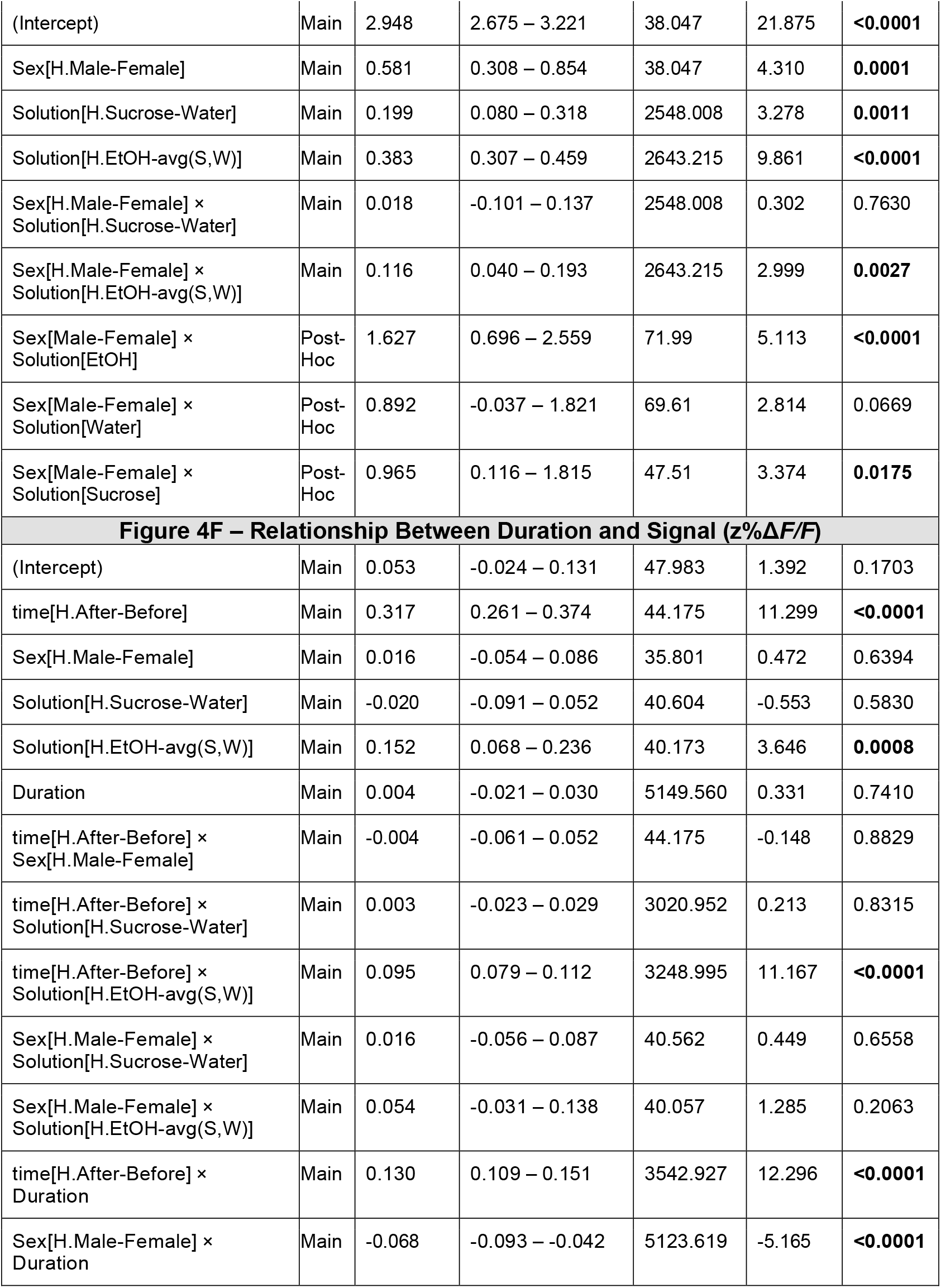
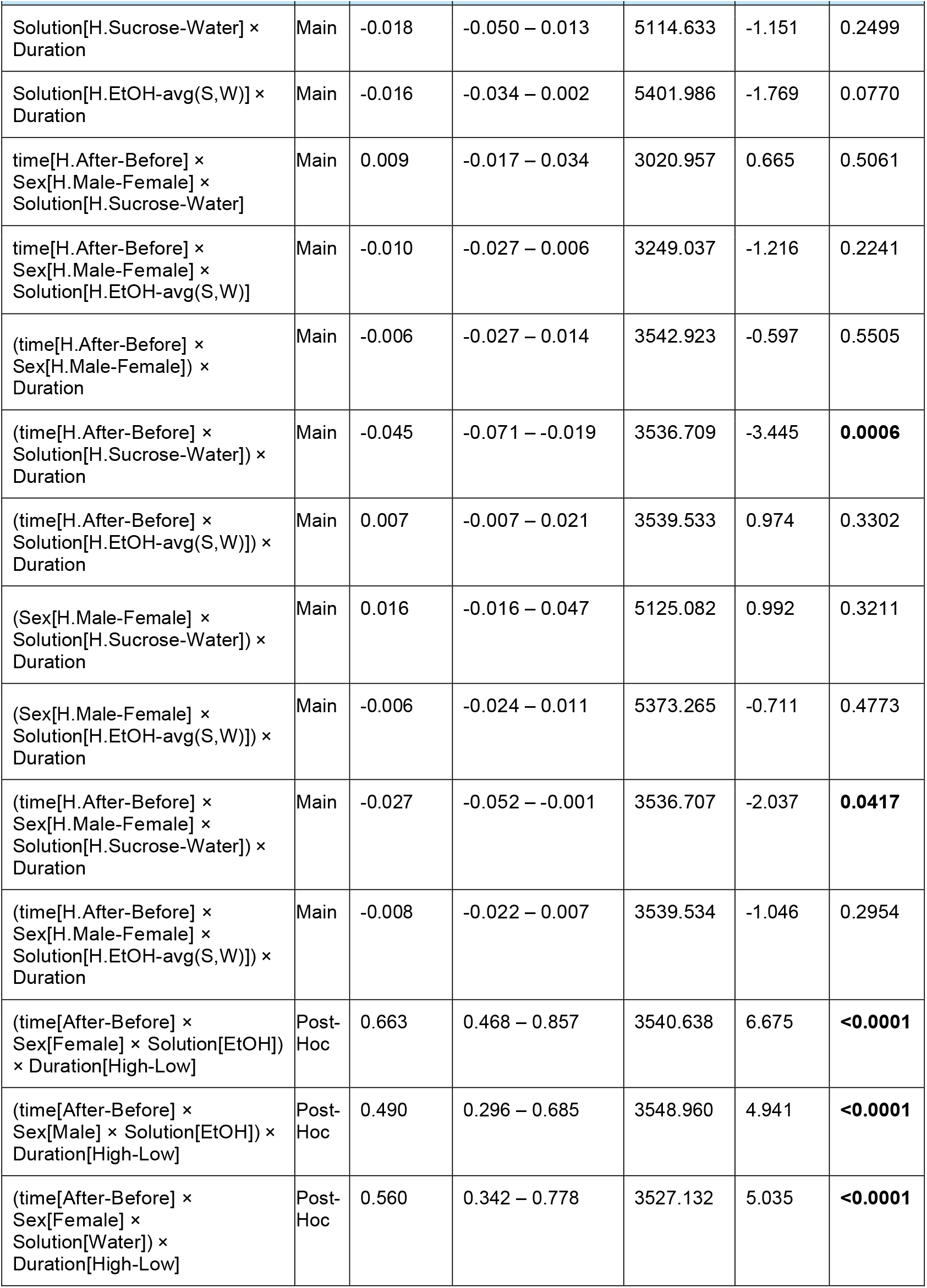
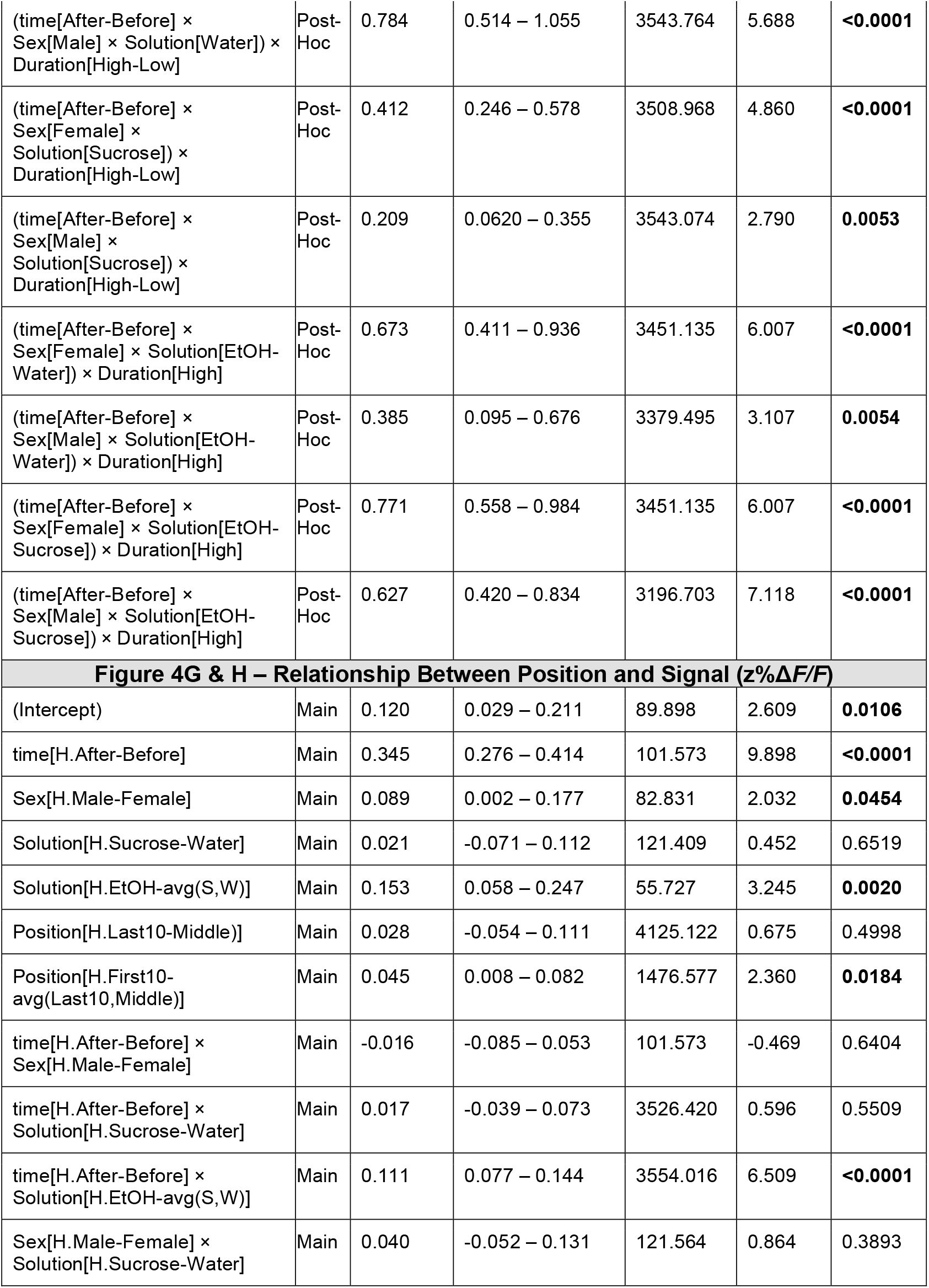
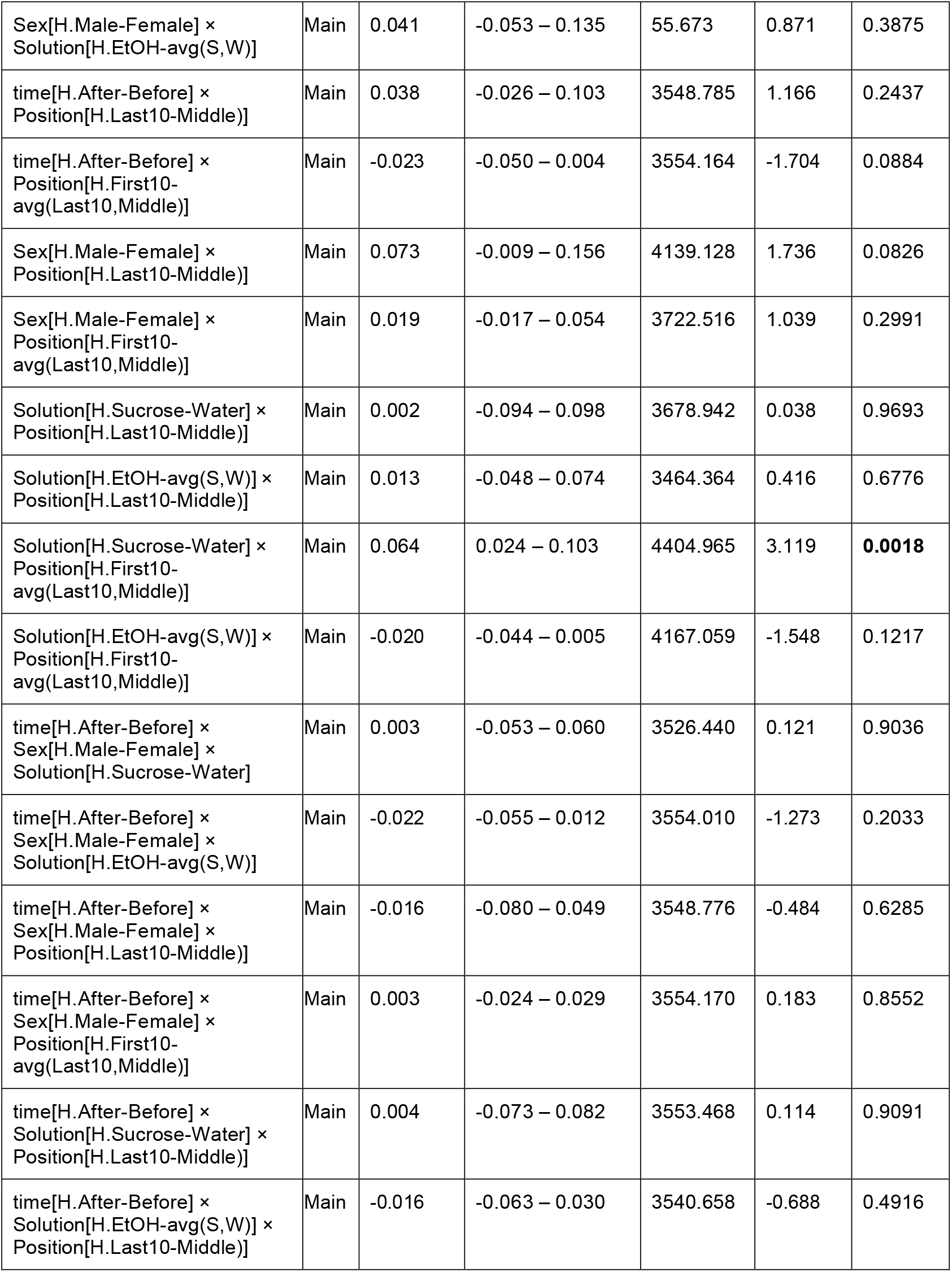
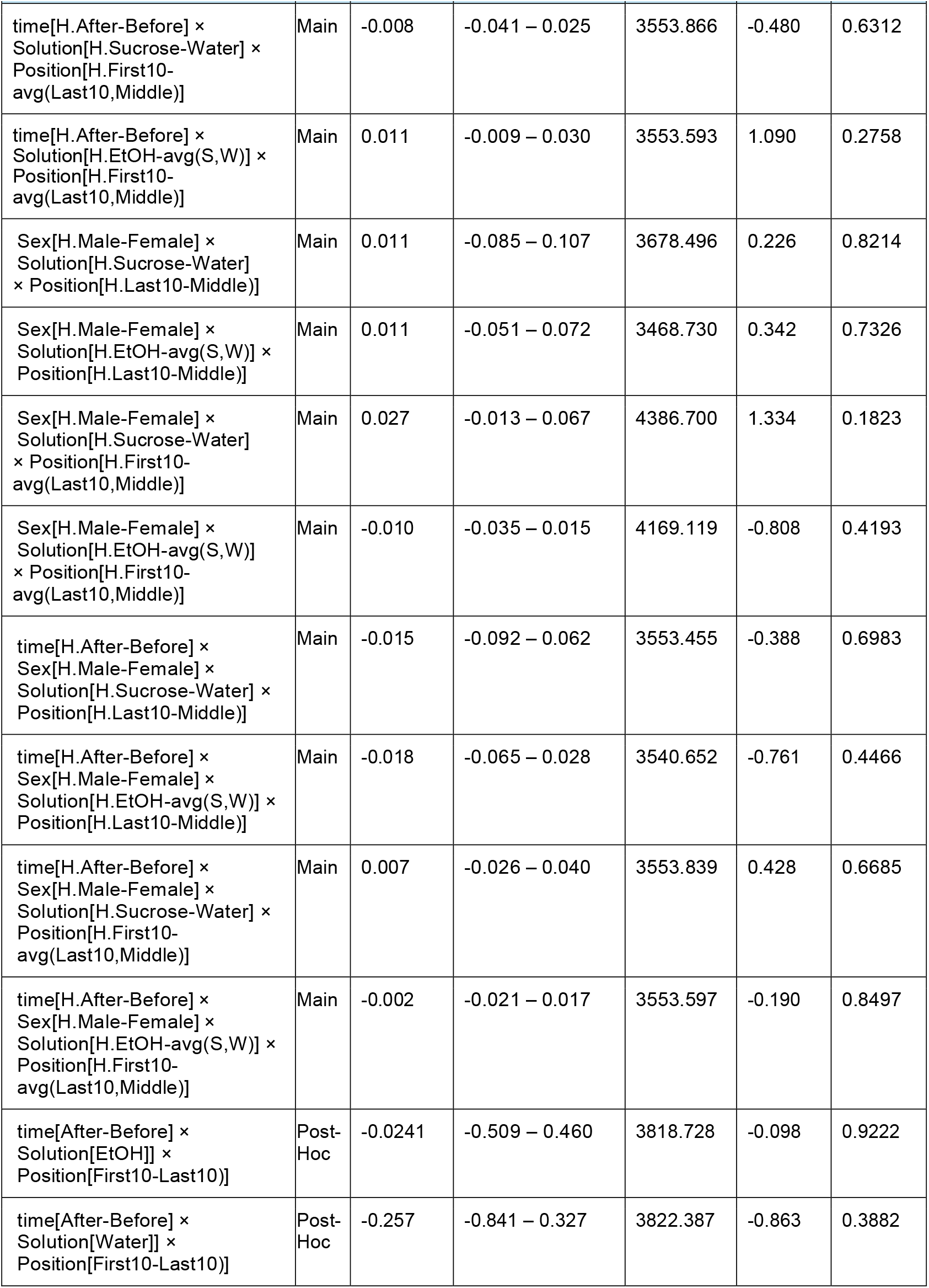
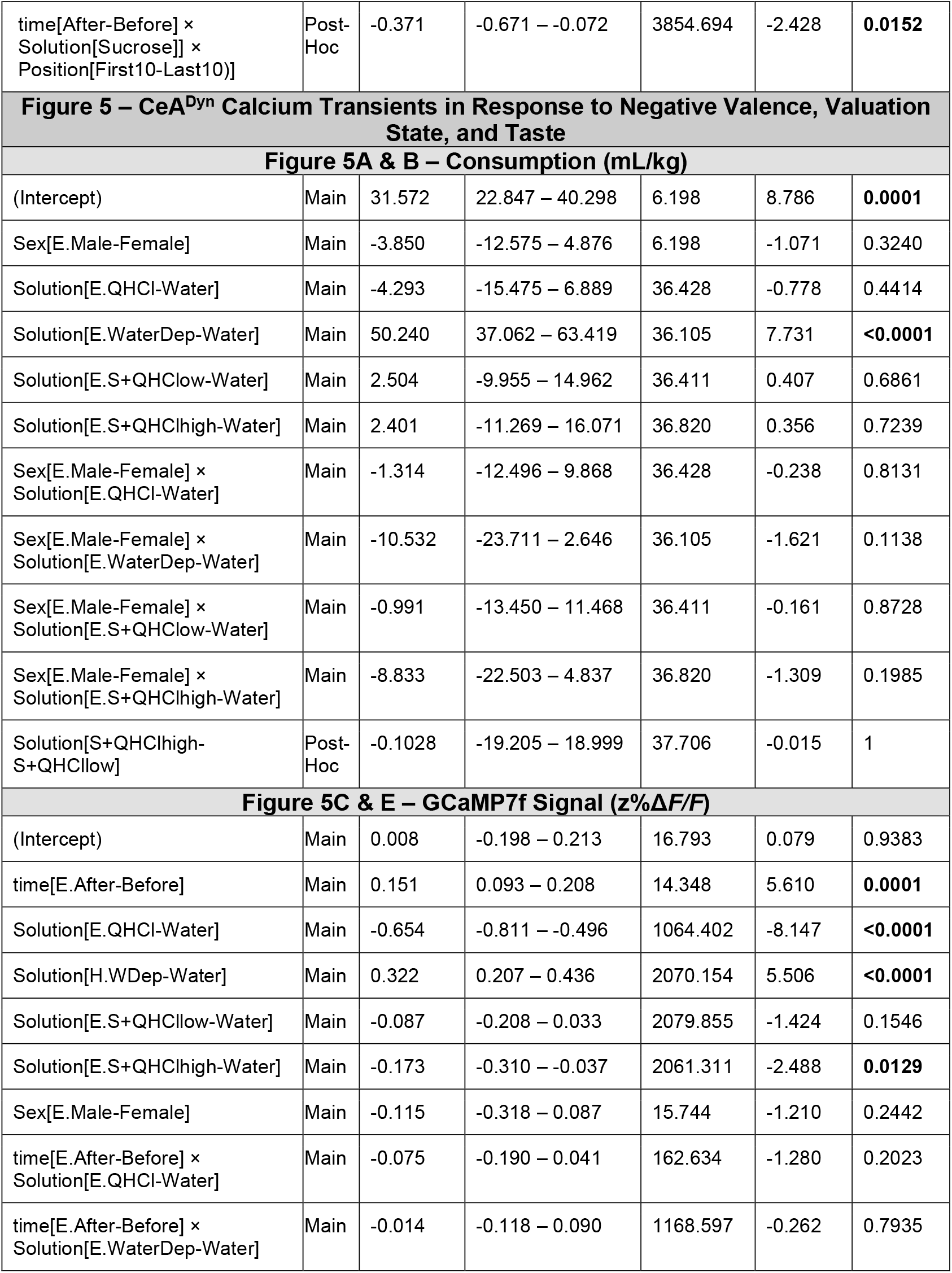
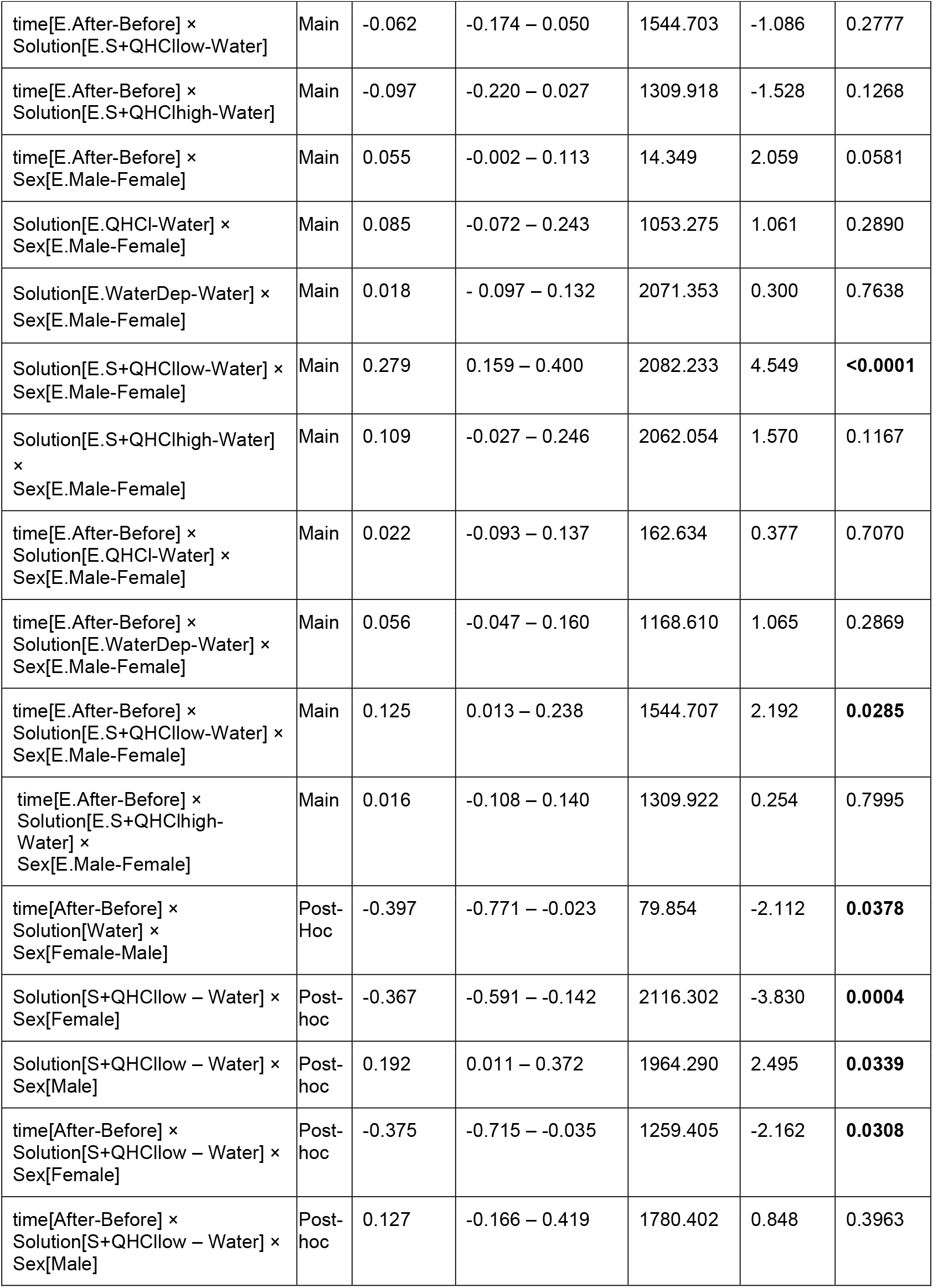
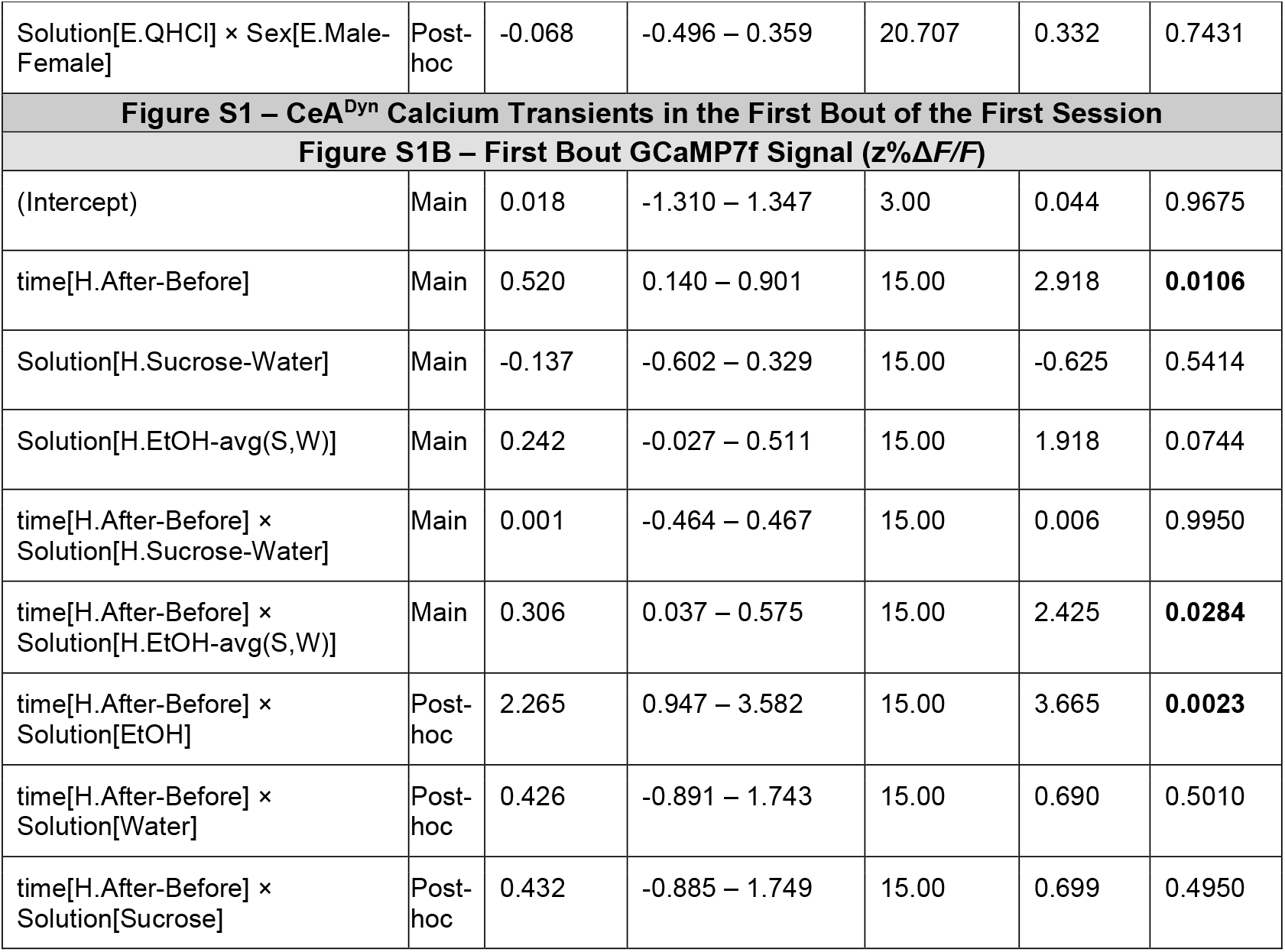

| Contrast<br>Factor [Contrast] x Interaction | Type | Estimate | 95% CI | df | Statistic | p Value |
| --- | --- | --- | --- | --- | --- | --- |
| **Figure 2 – Photometry Control Signal Correction** |  |  |  |  |  |  |
| **Figure 2C – Average 470 nm Signal During 405 nm Deflections (%$\Delta F/F$)** |  |  |  |  |  |  |
| (Intercept) | Main | 5.132 | 3.064 – 7.199 | 11.721 | 5.423 | **0.0002** |
| Method[E.Standard] | Main | 3.870 | 3.391 – 4.350 | 1531.821 | 15.840 | **<0.0001** |
| Method[E.Robust] | Main | -2.592 | -3.071 – -2.112 | 1531.821 | -10.607 | **<0.0001** |
| Method[E.Joint] | Main | 0.101 | -0.378 – 0.580 | 1531.821 | 0.413 | 0.6794 |
| Sex[H.Male-Female] | Main | 0.647 | -1.420 – 2.714 | 11.721 | 0.684 | 0.5076 |
| Solution[H.Sucrose-Water] | Main | -0.478 | -0.690 – -0.266 | 1532.472 | -4.415 | **<0.0001** |
| Solution[H.EtOH-Avg(S,W)] | Main | 0.261 | 0.136 – 0.385 | 1538.490 | 4.118 | **<0.0001** |
| Method[E.Standard] × Sex[H.Male-Female] | Main | 0.262 | -0.218 – 0.741 | 1531.821 | 1.071 | 0.2842 |
| Method[E.Robust] × Sex[H.Male-Female] | Main | -0.951 | -1.430 – -0.472 | 1531.821 | -3.891 | **0.0001** |
| Method[E.Joint] × Sex[H.Male-Female] | Main | 0.019 | -0.460 – 0.499 | 1531.821 | 0.079 | 0.9369 |
| Method[E.Standard] × Solution[H.Sucrose-Water] | Main | 0.009 | -0.582 – 0.601 | 1531.821 | 0.031 | 0.9749 |
| Method[E.Robust] × Solution[H.Sucrose-Water] | Main | 0.736 | 0.144 – 1.328 | 1531.821 | 2.438 | **0.0149** |
| Method[E.Joint] × Solution[H.Sucrose-Water] | Main | -0.002 | -0.594 – 0.590 | 1531.821 | -0.007 | 0.9943 |
| Method[E.Standard] × Solution[H.EtOH-Avg(S,W)] | Main | 0.014 | -0.322 – 0.350 | 1531.821 | 0.084 | 0.9331 |
| Method[E.Robust] × Solution[H.EtOH-Avg(S,W)] | Main | -0.039 | -0.375 – 0.297 | 1531.821 | -0.230 | 0.8180 |
| Method[E.Joint] × Solution[H.EtOH-Avg(S,W)] | Main | -0.001 | -0.337 – 0.335 | 1531.821 | -0.004 | 0.9964 |
| Sex[H.Male-Female] × Solution[H.Sucrose-Water] | Main | -0.251 | -0.463 – -0.038 | 1532.472 | -2.316 | **0.0207** |
| Sex[H.Male-Female] × Solution[H.EtOH-Avg(S,W)] | Main | 0.318 | 0.194 – 0.442 | 1538.490 | 5.022 | **<0.0001** |
| Method[E.Standard] × Sex[H.Male-Female] × Solution[H.Sucrose-Water] | Main | -0.044 | -0.636 – 0.548 | 1531.821 | -0.145 | 0.8847 |

| **Contrast**<br>Factor [Contrast] x Interaction | **Type** | **Estimate** | **95% CI** | **df** | **Statistic** | **p Value** |
| --- | --- | --- | --- | --- | --- | --- |
| Method[E.Robust] × Sex[H.Male-Female] × Solution[H.Sucrose-Water] | Main | 0.529 | -0.063 – 1.121 | 1531.821 | 1.754 | 0.0796 |
| Method[E.Joint] × Sex[H.Male-Female] × Solution[H.Sucrose-Water] | Main | 0.000 | -0.592 – 0.592 | 1531.821 | 0.001 | 0.9993 |
| Method[E.Standard] × Sex[H.Male-Female] × Solution[H.EtOH-Avg(S,W)] | Main | 0.081 | -0.255 – 0.417 | 1531.821 | 0.473 | 0.6365 |
| Method[E.Robust] × Sex[H.Male-Female] × Solution[H.EtOH-Avg(S,W)] | Main | 0.257 | -0.079 – 0.593 | 1531.821 | 1.502 | 0.1333 |
| Method[E.Joint] × Sex[H.Male-Female] × Solution[H.EtOH-Avg(S,W)] | Main | 0.002 | -0.334 – 0.338 | 1531.821 | 0.010 | 0.9917 |
| **Figure 2D – Average 470 nm SignalMAD** |  |  |  |  |  |  |
| (Intercept) | Main | 1.622 | 1.241 – 2.003 | 13.012 | 9.201 | **<0.0001** |
| Method[E.Standard] | Main | 0.259 | 0.189 – 0.330 | 472.174 | 7.224 | **<0.0001** |
| Method[E.Robust] | Main | -0.019 | -0.089 – 0.052 | 472.174 | -0.518 | 0.6049 |
| Method[E.Joint] | Main | 0.062 | -0.008 – 0.133 | 472.174 | 1.729 | 0.0845 |
| Solution[H.Sucrose-Water] | Main | -0.028 | -0.061 – 0.005 | 515.025 | -1.687 | 0.0922 |
| Solution[H.EtOH-avg(S,W)] | Main | 0.050 | 0.032 – 0.067 | 513.209 | 5.505 | **<0.0001** |
| Method[E.Standard] × Solution[H.Sucrose-Water] | Main | -0.008 | -0.097 – 0.081 | 472.174 | -0.175 | 0.8608 |
| Method[E.Robust] × Solution[H.Sucrose-Water] | Main | 0.011 | -0.078 – 0.100 | 472.174 | 0.241 | 0.8093 |
| Method[E.Joint] × Solution[H.Sucrose-Water] | Main | 0.002 | -0.087 – 0.091 | 472.174 | 0.041 | 0.9673 |
| Method[E.Standard] × Solution[H.EtOH-avg(S,W)] | Main | 0.014 | -0.035 – 0.062 | 472.174 | 0.558 | 0.5770 |
| Method[E.Robust] × Solution[H.EtOH-avg(S,W)] | Main | -0.031 | -0.080 – 0.017 | 472.174 | -1.277 | 0.2021 |
| Method[E.Joint] × Solution[H.EtOH-avg(S,W)] | Main | 0.003 | -0.046 – 0.051 | 472.174 | 0.106 | 0.9158 |
| **Figure 2E – Average 470 nm SignalMean Peak Frequency (Peaks/Second)** |  |  |  |  |  |  |
| (Intercept) | Main | 0.137 | 0.103 – 0.171 | 13.056 | 8.711 | **<0.0001** |
| Method[E.Standard] | Main | -0.031 | -0.038 – -0.023 | 466.988 | -8.050 | **<0.0001** |
| Method[E.Robust] | Main | 0.023 | 0.016 – 0.031 | 466.988 | 6.153 | **<0.0001** |
| Method[E.Joint] | Main | 0.007 | -0.000 – 0.015 | 466.988 | 1.891 | 0.0593 |
| Robust - Joint | Post<br>-Hoc | 0.0163 | 0.006 – 0.026 | 466.988 | 4.262 | **0.0001** |
| **Figure 2F – Average 470 nm SignalMean Peak Amplitude (%$\Delta F/F$)** |  |  |  |  |  |  |
| (Intercept) | Main | 4.270 | 3.091 – 5.449 | 13.003 | 7.826 | **<0.0001** |
| Method[E.Standard] | Main | 0.589 | 0.420 – 0.757 | 472.659 | 6.871 | **<0.0001** |
| Method[E.Robust] | Main | -0.189 | -0.357 – -0.021 | 472.659 | -2.205 | **0.0279** |
| Method[E.Joint] | Main | 0.250 | 0.082 – 0.418 | 472.659 | 2.918 | **0.0037** |
| Solution[H.Sucrose-Water] | Main | -0.121 | -0.201 – -0.041 | 511.107 | -2.970 | **0.0031** |
| Solution[H.EtOH-avg(S,W)] | Main | 0.209 | 0.166 – 0.251 | 508.392 | 9.596 | **<0.0001** |
| Method[E.Standard] ×<br>Solution[H.Sucrose-Water] | Main | 0.007 | -0.206 – 0.220 | 472.659 | 0.063 | 0.9496 |
| Method[E.Robust] ×<br>Solution[H.Sucrose-Water] | Main | 0.034 | -0.179 – 0.247 | 472.659 | 0.315 | 0.7532 |
| Method[E.Joint] ×<br>Solution[H.Sucrose-Water] | Main | -0.000 | -0.213 – 0.213 | 472.659 | -0.001 | 0.9989 |
| Method[E.Standard] ×<br>Solution[H.EtOH-avg(S,W)] | Main | 0.011 | -0.104 – 0.126 | 472.659 | 0.191 | 0.8489 |
| Method[E.Robust] ×<br>Solution[H.EtOH-avg(S,W)] | Main | -0.066 | -0.181 – 0.049 | 472.659 | -1.128 | 0.2599 |
| Method[E.Joint] ×<br>Solution[H.EtOH-avg(S,W)] | Main | 0.006 | -0.109 – 0.121 | 472.659 | 0.108 | 0.9140 |
| Standard – Joint | Post<br>-Hoc | 0.339 | 0.118 – 0.559 | 472.659 | 3.953 | **0.0005** |
| **Figure 2H – 470 nm Signal Slope DuringMovement Artifact (%$\Delta F/F$ / Seconds)** |  |  |  |  |  |  |
| (Intercept) | Main | -0.183 | -1.017 – 0.651 | 7.966 | -0.506 | 0.6264 |
| Method[E.Standard] | Main | 9.042 | 7.497 – 10.586 | 94.622 | 11.622 | **<0.0001** |
| Method[E.Robust] | Main | 4.960 | 3.416 – 6.505 | 94.622 | 6.376 | **<0.0001** |
| Method[E.Joint] | Main | 8.820 | 7.275 – 10.364 | 94.622 | 11.337 | **<0.0001** |
| Sex[H.Male-Female] | Main | -0.672 | -1.506 – 0.162 | 7.966 | -1.860 | 0.1001 |
| Method[E.Standard] ×<br>Sex[H.Male-Female] | Main | -1.884 | -3.429 – -0.340 | 94.622 | -2.422 | **0.0173** |
| Method[E.Robust] ×<br>Sex[H.Male-Female] | Main | -1.372 | -2.916 – 0.173 | 94.622 | -1.763 | 0.0811 |

| **Contrast**<br>Factor [Contrast] x Interaction | **Type** | **Estimate** | **95% CI** | **df** | **Statistic** | **p Value** |
| --- | --- | --- | --- | --- | --- | --- |
| Method[E.Joint] × Sex[H.Male-Female] | Main | -2.103 | -3.647 – -0.558 | 94.622 | -2.703 | **0.0082** |
| Method[Standard – Robust] | Post-Hoc | 4.081 | 2.047 – 6.116 | 94.622 | 5.246 | **<0.0001** |
| Method[Standard – Joint] | Post-Hoc | 0.222 | -1.813 – 2.256 | 94.622 | 0.285 | 0.9919 |
| Method[Robust – Joint] | Post-Hoc | -3.860 | -5.894 – -1.825 | 94.622 | -4.961 | **<0.0001** |
| **Figure 2I – 470 nm Signal Correlation (With 405 nm Signal)** |  |  |  |  |  |  |
| (Intercept) | Main | -0.169 | -0.397 – 0.059 | 6.992 | -1.753 | 0.1230 |
| Method[E.Standard] | Main | -1.894 | -2.154 – -1.634 | 97.103 | -14.461 | **<0.0001** |
| Method[E.Robust] | Main | -0.890 | -1.150 – -0.630 | 97.103 | -6.796 | **<0.0001** |
| Method[E.Joint] | Main | -1.883 | -2.143 – -1.623 | 97.103 | -14.372 | **<0.0001** |
| Method[Standard – Robust] | Post-Hoc | -1.004 | -1.347 – -0.662 | 97.103 | -7.665 | **<0.0001** |
| Method[Standard – Joint] | Post-Hoc | -0.012 | -0.354 – 0.331 | 97.103 | -0.089 | 0.9997 |
| Method[Robust – Joint] | Post-Hoc | 0.992 | 0.650 – 1.335 | 97.103 | 7.575 | **<0.0001** |
| **Figure 3B & D – GCaMP7f Signal ( $z\%\Delta F/F$ )** |  |  |  |  |  |  |
| (Intercept) | Main | 0.052 | -0.027 – 0.130 | 46.615 | 1.325 | 0.1916 |
| time[H.After – Before] | Main | 0.327 | 0.271 – 0.383 | 43.533 | 11.747 | **<0.0001** |
| Sex[H.Male – Female] | Main | 0.012 | -0.060 – 0.083 | 35.081 | 0.331 | 0.7430 |
| Solution[H.Sucrose – Water] | Main | -0.016 | -0.088 – 0.056 | 39.516 | -0.448 | 0.6563 |
| Solution[H.EtOH – avg(S,W)] | Main | 0.152 | 0.067 – 0.236 | 39.888 | 3.611 | **0.0008** |
| time[H.After – Before] × Sex[H.Male – Female] | Main | -0.004 | -0.060 – 0.052 | 43.533 | -0.157 | 0.8759 |
| time[H.After – Before] × Solution[H.Sucrose – Water] | Main | 0.020 | -0.006 – 0.046 | 3001.772 | 1.541 | 0.1233 |
| time[H.After – Before] × Solution[H.EtOH – avg(S,W)] | Main | 0.118 | 0.102 – 0.135 | 3160.989 | 14.060 | **<0.0001** |
| Sex[H.Male – Female] × Solution[H.Sucrose – Water] | Main | 0.007 | -0.065 – 0.079 | 39.473 | 0.199 | 0.8436 |

| **Contrast**<br>Factor [Contrast] x Interaction | **Type** | **Estimate** | **95% CI** | **df** | **Statistic** | **p Value** |
| --- | --- | --- | --- | --- | --- | --- |
| Sex[H.Male – Female] ×<br>Solution[H.EtOH – avg(S,W)] | Main | 0.040 | -0.045 – 0.125 | 39.798 | 0.953 | 0.3465 |
| time[H.After – Before] ×<br>Sex[H.Male – Female] ×<br>Solution[H.Sucrose – Water] | Main | 0.012 | -0.014 – 0.038 | 3001.795 | 0.918 | 0.3588 |
| time[H.After – Before] ×<br>Sex[H.Male – Female] ×<br>Solution[H.EtOH – avg(S,W)] | Main | -0.009 | -0.026 – 0.007 | 3161.023 | -1.093 | 0.2745 |
| time[After – Before] ×<br>Solution[EtOH – Water] | Post-Hoc | 0.750 | 0.607 – 0.893 | 3075.36 | 12.348 | **<0.0001** |
| time[After – Before] ×<br>Solution[EtOH – Sucrose] | Post-Hoc | 0.669 | 0.545 – 0.793 | 3194.74 | 12.673 | **<0.0001** |
| Time[After – Before] x<br>Sex[H.Male – Female] x<br>Solution[Water] | Post-Hoc | -0.029 | -0.293 – 0.235 | 83.21 | -0.219 | 0.8275 |
| **Figure 4A & B – Consumption (mL/kg)** |  |  |  |  |  |  |
| (Intercept) | Main | 23.972 | 20.428 – 27.517 | 12.096 | 14.722 | **<0.0001** |
| Sex[H.Male – Female] | Main | -4.998 | -8.543 – -1.453 | 12.096 | -3.069 | **0.0096** |
| Solution[H.Sucrose – Water] | Main | 3.197 | 0.956 – 5.439 | 104.716 | 2.828 | **0.0056** |
| Solution[H.EtOH – avg(S,W)] | Main | -4.401 | -5.649 – -3.152 | 105.651 | -6.986 | **<0.0001** |
| Sex[H.Male – Female] ×<br>Solution[H.Sucrose – Water] | Main | 0.539 | -1.702 – 2.781 | 104.716 | 0.477 | 0.6342 |
| Sex[H.Male – Female] ×<br>Solution[H.EtOH – avg(S,W)] | Main | 1.671 | 0.422 – 2.920 | 105.651 | 2.652 | **0.0092** |
| Sex[Female – Male] ×<br>Solution[EtOH] | Post-Hoc | 3.313 | -9.106 – 15.732 | 27.85 | 0.815 | 0.9622 |
| Sex[Female – Male] ×<br>Solution[Water] | Post-Hoc | 14.416 | 1.131 – 27.700 | 38.08 | 3.255 | **0.0267** |
| Sex[Female – Male] ×<br>Solution[Sucrose] | Post-Hoc | 12.258 | 0.145 – 24.372 | 24.96 | 3.119 | **0.0461** |
| Sex[Female] x Solution[Water -<br>Sucrose] | Post-Hoc | -5.316 | -15.166 – 4.534 | 104.916 | -1.566 | 0.6229 |
| Sex[Female] x Solution[Water -<br>EtOH] | Post-Hoc | 15.556 | 5.474 – 25.637 | 105.147 | 4.475 | **0.0003** |
| Sex[Female] x Solution[Sucrose<br>- EtOH] | Post-Hoc | 20.872 | 11.610 – 30.134 | 106.589 | 6.525 | **<0.0001** |
| Sex[Male] x Solution[Water - Sucrose] | Post-Hoc | -7.474 | -16.152 – 1.204 | 104.600 | -2.499 | 0.1338 |
| Sex[Male] x Solution[Water - EtOH] | Post-Hoc | 4.452 | -4.415 – 13.319 | 104.532 | 1.457 | 0.6920 |
| Sex[Male] x Solution[Sucrose - EtOH] | Post-Hoc | 11.926 | 4.126 – 19.726 | 105.494 | 4.433 | **0.0003** |
| **Figure 4C – Relationship Between Consumption and Signal ($z\%\Delta F/F$)** |  |  |  |  |  |  |
| (Intercept) | Main | 0.041 | -0.080 – 0.161 | 40.166 | 0.685 | 0.4970 |
| time[H.After-Before] | Main | 0.310 | 0.251 – 0.370 | 52.247 | 10.449 | **<0.0001** |
| Sex[H.Male-Female] | Main | -0.092 | -0.209 – 0.024 | 34.808 | -1.611 | 0.1162 |
| Solution[H.Sucrose-Water] | Main | -0.004 | -0.085 – 0.076 | 37.293 | -0.104 | 0.9175 |
| Solution[H.EtOH-avg(S,W)] | Main | 0.139 | 0.017 – 0.261 | 40.081 | 2.308 | **0.0262** |
| Consumption | Main | -0.025 | -0.116 – 0.066 | 65.061 | -0.558 | 0.5788 |
| time[H.After-Before] × Sex[H.Male-Female] | Main | -0.046 | -0.105 – 0.014 | 52.247 | -1.537 | 0.1303 |
| time[H.After-Before] × Solution[H.Sucrose-Water] | Main | 0.033 | 0.004 – 0.062 | 2623.325 | 2.227 | **0.0260** |
| time[H.After-Before] × Solution[H.EtOH-avg(S,W)] | Main | 0.115 | 0.090 – 0.140 | 2386.977 | 9.131 | **<0.0001** |
| Sex[H.Male-Female] × Solution[H.Sucrose-Water] | Main | 0.020 | -0.060 – 0.101 | 37.293 | 0.507 | 0.6148 |
| Sex[H.Male-Female] × Solution[H.EtOH-avg(S,W)] | Main | -0.050 | -0.172 – 0.072 | 40.079 | -0.822 | 0.4157 |
| time[H.After-Before] × Consumption | Main | -0.037 | -0.065 – -0.009 | 2255.628 | -2.569 | **0.0103** |
| Sex[H.Male-Female] × Consumption | Main | -0.095 | -0.186 – -0.005 | 64.613 | -2.099 | **0.0397** |
| Solution[H.Sucrose-Water] × Consumption | Main | -0.019 | -0.090 – 0.053 | 54.653 | -0.525 | 0.6016 |
| Solution[H.EtOH-avg(S,W)] × Consumption | Main | -0.003 | -0.077 – 0.072 | 33.220 | -0.073 | 0.9419 |
| time[H.After-Before] × Sex[H.Male-Female] × Solution[H.Sucrose-Water] | Main | 0.019 | -0.010 – 0.048 | 2623.340 | 1.314 | 0.1889 |
| time[H.After-Before] × Sex[H.Male-Female] × Solution[H.EtOH-avg(S,W)] | Main | -0.035 | -0.060 – -0.011 | 2387.011 | -2.804 | **0.0051** |

| **Contrast**<br>Factor [Contrast] x Interaction | **Type** | **Estimate** | **95% CI** | **df** | **Statistic** | **p Value** |
| --- | --- | --- | --- | --- | --- | --- |
| (time[H.After-Before] × Sex[H.Male-Female]) × Consumption | Main | -0.045 | -0.073 – -0.017 | 2255.667 | -3.114 | **0.0019** |
| (time[H.After-Before] × Solution[H.Sucrose-Water]) × Consumption | Main | 0.034 | -0.003 – 0.070 | 1308.916 | 1.800 | 0.0720 |
| (time[H.After-Before] × Solution[H.EtOH-avg(S,W)]) × Consumption | Main | 0.007 | -0.016 – 0.030 | 1613.016 | 0.579 | 0.5626 |
| (Sex[H.Male-Female] × Solution[H.Sucrose-Water]) × Consumption | Main | 0.035 | -0.037 – 0.106 | 54.612 | 0.979 | 0.3320 |
| (Sex[H.Male-Female] × Solution[H.EtOH-avg(S,W)]) × Consumption | Main | -0.065 | -0.140 – 0.009 | 33.174 | -1.784 | 0.0836 |
| (time[H.After-Before] × Sex[H.Male-Female] × Solution[H.Sucrose-Water]) × Consumption | Main | 0.030 | -0.007 – 0.066 | 1308.909 | 1.593 | 0.1113 |
| (time[H.After-Before] × Sex[H.Male-Female] × Solution[H.EtOH-avg(S,W)]) × Consumption | Main | -0.027 | -0.050 – -0.004 | 1612.916 | -2.259 | **0.0240** |
| Time[After-Before] × Sex[Female] × Solution[EtOH] × Drinking[High-Low] | Post-Hoc | 0.301 | -0.041 – 0.643 | 2308.937 | 1.729 | 0.08402 |
| Time[After-Before] × Sex[Male] × Solution[EtOH] × Drinking[High-Low] | Post-Hoc | -0.486 | -0.761 – -0.211 | 2693.910 | -3.469 | **0.0005** |
| Time[After-Before] × Sex[Female] × Solution[Water] × Drinking[High-Low] | Post-Hoc | -0.119 | -0.380 – 0.141 | 2795.112 | -0.899 | 0.3688 |
| Time[After-Before] × Sex[Male] × Solution[Water] × Drinking[High-Low] | Post-Hoc | -0.499 | -0.817 – -0.181 | 1525.602 | -3.074 | **0.0021** |
| Time[After-Before] × Sex[Female] × Solution[Sucrose] × Drinking[High-Low] | Post-Hoc | -0.088 | -0.384 – 0.207 | 905.577 | -0.587 | 0.5571 |
| Time[After-Before] × Sex[Male] × Solution[Sucrose] × Drinking[High-Low] | Post-Hoc | 0.007 | -0.282 – 0.295 | 901.718 | 0.0466 | 0.9629 |
| **Figure 4D & E – Bout Duration (seconds)** |  |  |  |  |  |  |
| (Intercept) | Main | 2.948 | 2.675 – 3.221 | 38.047 | 21.875 | **<0.0001** |
| Sex[H.Male-Female] | Main | 0.581 | 0.308 – 0.854 | 38.047 | 4.310 | **0.0001** |
| Solution[H.Sucrose-Water] | Main | 0.199 | 0.080 – 0.318 | 2548.008 | 3.278 | **0.0011** |
| Solution[H.EtOH-avg(S,W)] | Main | 0.383 | 0.307 – 0.459 | 2643.215 | 9.861 | **<0.0001** |
| Sex[H.Male-Female] ×<br>Solution[H.Sucrose-Water] | Main | 0.018 | -0.101 – 0.137 | 2548.008 | 0.302 | 0.7630 |
| Sex[H.Male-Female] ×<br>Solution[H.EtOH-avg(S,W)] | Main | 0.116 | 0.040 – 0.193 | 2643.215 | 2.999 | **0.0027** |
| Sex[Male-Female] ×<br>Solution[EtOH] | Post-Hoc | 1.627 | 0.696 – 2.559 | 71.99 | 5.113 | **<0.0001** |
| Sex[Male-Female] ×<br>Solution[Water] | Post-Hoc | 0.892 | -0.037 – 1.821 | 69.61 | 2.814 | 0.0669 |
| Sex[Male-Female] ×<br>Solution[Sucrose] | Post-Hoc | 0.965 | 0.116 – 1.815 | 47.51 | 3.374 | **0.0175** |
| **Figure 4F – Relationship Between Duration and Signal ( $z\%\Delta F/F$ )** |  |  |  |  |  |  |
| (Intercept) | Main | 0.053 | -0.024 – 0.131 | 47.983 | 1.392 | 0.1703 |
| time[H.After-Before] | Main | 0.317 | 0.261 – 0.374 | 44.175 | 11.299 | **<0.0001** |
| Sex[H.Male-Female] | Main | 0.016 | -0.054 – 0.086 | 35.801 | 0.472 | 0.6394 |
| Solution[H.Sucrose-Water] | Main | -0.020 | -0.091 – 0.052 | 40.604 | -0.553 | 0.5830 |
| Solution[H.EtOH-avg(S,W)] | Main | 0.152 | 0.068 – 0.236 | 40.173 | 3.646 | **0.0008** |
| Duration | Main | 0.004 | -0.021 – 0.030 | 5149.560 | 0.331 | 0.7410 |
| time[H.After-Before] ×<br>Sex[H.Male-Female] | Main | -0.004 | -0.061 – 0.052 | 44.175 | -0.148 | 0.8829 |
| time[H.After-Before] ×<br>Solution[H.Sucrose-Water] | Main | 0.003 | -0.023 – 0.029 | 3020.952 | 0.213 | 0.8315 |
| time[H.After-Before] ×<br>Solution[H.EtOH-avg(S,W)] | Main | 0.095 | 0.079 – 0.112 | 3248.995 | 11.167 | **<0.0001** |
| Sex[H.Male-Female] ×<br>Solution[H.Sucrose-Water] | Main | 0.016 | -0.056 – 0.087 | 40.562 | 0.449 | 0.6558 |
| Sex[H.Male-Female] ×<br>Solution[H.EtOH-avg(S,W)] | Main | 0.054 | -0.031 – 0.138 | 40.057 | 1.285 | 0.2063 |
| time[H.After-Before] ×<br>Duration | Main | 0.130 | 0.109 – 0.151 | 3542.927 | 12.296 | **<0.0001** |
| Sex[H.Male-Female] ×<br>Duration | Main | -0.068 | -0.093 – -0.042 | 5123.619 | -5.165 | **<0.0001** |

| **Contrast**<br>Factor [Contrast] x Interaction | **Type** | **Estimate** | **95% CI** | **df** | **Statistic** | **p Value** |
| --- | --- | --- | --- | --- | --- | --- |
| Solution[H.Sucrose-Water] × Duration | Main | -0.018 | -0.050 – 0.013 | 5114.633 | -1.151 | 0.2499 |
| Solution[H.EtOH-avg(S,W)] × Duration | Main | -0.016 | -0.034 – 0.002 | 5401.986 | -1.769 | 0.0770 |
| time[H.After-Before] × Sex[H.Male-Female] × Solution[H.Sucrose-Water] | Main | 0.009 | -0.017 – 0.034 | 3020.957 | 0.665 | 0.5061 |
| time[H.After-Before] × Sex[H.Male-Female] × Solution[H.EtOH-avg(S,W)] | Main | -0.010 | -0.027 – 0.006 | 3249.037 | -1.216 | 0.2241 |
| (time[H.After-Before] × Sex[H.Male-Female]) × Duration | Main | -0.006 | -0.027 – 0.014 | 3542.923 | -0.597 | 0.5505 |
| (time[H.After-Before] × Solution[H.Sucrose-Water]) × Duration | Main | -0.045 | -0.071 – -0.019 | 3536.709 | -3.445 | **0.0006** |
| (time[H.After-Before] × Solution[H.EtOH-avg(S,W)]) × Duration | Main | 0.007 | -0.007 – 0.021 | 3539.533 | 0.974 | 0.3302 |
| (Sex[H.Male-Female] × Solution[H.Sucrose-Water]) × Duration | Main | 0.016 | -0.016 – 0.047 | 5125.082 | 0.992 | 0.3211 |
| (Sex[H.Male-Female] × Solution[H.EtOH-avg(S,W)]) × Duration | Main | -0.006 | -0.024 – 0.011 | 5373.265 | -0.711 | 0.4773 |
| (time[H.After-Before] × Sex[H.Male-Female] × Solution[H.Sucrose-Water]) × Duration | Main | -0.027 | -0.052 – -0.001 | 3536.707 | -2.037 | **0.0417** |
| (time[H.After-Before] × Sex[H.Male-Female] × Solution[H.EtOH-avg(S,W)]) × Duration | Main | -0.008 | -0.022 – 0.007 | 3539.534 | -1.046 | 0.2954 |
| (time[After-Before] × Sex[Female] × Solution[EtOH]) × Duration[High-Low] | Post-Hoc | 0.663 | 0.468 – 0.857 | 3540.638 | 6.675 | **<0.0001** |
| (time[After-Before] × Sex[Male] × Solution[EtOH]) × Duration[High-Low] | Post-Hoc | 0.490 | 0.296 – 0.685 | 3548.960 | 4.941 | **<0.0001** |
| (time[After-Before] × Sex[Female] × Solution[Water]) × Duration[High-Low] | Post-Hoc | 0.560 | 0.342 – 0.778 | 3527.132 | 5.035 | **<0.0001** |
| (time[After-Before] × Sex[Male] × Solution[Water]) × Duration[High-Low] | Post-Hoc | 0.784 | 0.514 – 1.055 | 3543.764 | 5.688 | **<0.0001** |
| (time[After-Before] × Sex[Female] × Solution[Sucrose]) × Duration[High-Low] | Post-Hoc | 0.412 | 0.246 – 0.578 | 3508.968 | 4.860 | **<0.0001** |
| (time[After-Before] × Sex[Male] × Solution[Sucrose]) × Duration[High-Low] | Post-Hoc | 0.209 | 0.0620 – 0.355 | 3543.074 | 2.790 | **0.0053** |
| (time[After-Before] × Sex[Female] × Solution[EtOH-Water]) × Duration[High] | Post-Hoc | 0.673 | 0.411 – 0.936 | 3451.135 | 6.007 | **<0.0001** |
| (time[After-Before] × Sex[Male] × Solution[EtOH-Water]) × Duration[High] | Post-Hoc | 0.385 | 0.095 – 0.676 | 3379.495 | 3.107 | **0.0054** |
| (time[After-Before] × Sex[Female] × Solution[EtOH-Sucrose]) × Duration[High] | Post-Hoc | 0.771 | 0.558 – 0.984 | 3451.135 | 6.007 | **<0.0001** |
| (time[After-Before] × Sex[Male] × Solution[EtOH-Sucrose]) × Duration[High] | Post-Hoc | 0.627 | 0.420 – 0.834 | 3196.703 | 7.118 | **<0.0001** |
| **Figure 4G & H – Relationship Between Position and Signal (z%ΔF/F)** |  |  |  |  |  |  |
| (Intercept) | Main | 0.120 | 0.029 – 0.211 | 89.898 | 2.609 | **0.0106** |
| time[H.After-Before] | Main | 0.345 | 0.276 – 0.414 | 101.573 | 9.898 | **<0.0001** |
| Sex[H.Male-Female] | Main | 0.089 | 0.002 – 0.177 | 82.831 | 2.032 | **0.0454** |
| Solution[H.Sucrose-Water] | Main | 0.021 | -0.071 – 0.112 | 121.409 | 0.452 | 0.6519 |
| Solution[H.EtOH-avg(S,W)] | Main | 0.153 | 0.058 – 0.247 | 55.727 | 3.245 | **0.0020** |
| Position[H.Last10-Middle]) | Main | 0.028 | -0.054 – 0.111 | 4125.122 | 0.675 | 0.4998 |
| Position[H.First10-avg(Last10,Middle)] | Main | 0.045 | 0.008 – 0.082 | 1476.577 | 2.360 | **0.0184** |
| time[H.After-Before] × Sex[H.Male-Female] | Main | -0.016 | -0.085 – 0.053 | 101.573 | -0.469 | 0.6404 |
| time[H.After-Before] × Solution[H.Sucrose-Water] | Main | 0.017 | -0.039 – 0.073 | 3526.420 | 0.596 | 0.5509 |
| time[H.After-Before] × Solution[H.EtOH-avg(S,W)] | Main | 0.111 | 0.077 – 0.144 | 3554.016 | 6.509 | **<0.0001** |
| Sex[H.Male-Female] × Solution[H.Sucrose-Water] | Main | 0.040 | -0.052 – 0.131 | 121.564 | 0.864 | 0.3893 |
| Sex[H.Male-Female] ×<br>Solution[H.EtOH-avg(S,W)] | Main | 0.041 | -0.053 – 0.135 | 55.673 | 0.871 | 0.3875 |
| time[H.After-Before] ×<br>Position[H.Last10-Middle]] | Main | 0.038 | -0.026 – 0.103 | 3548.785 | 1.166 | 0.2437 |
| time[H.After-Before] ×<br>Position[H.First10-avg(Last10,Middle)] | Main | -0.023 | -0.050 – 0.004 | 3554.164 | -1.704 | 0.0884 |
| Sex[H.Male-Female] ×<br>Position[H.Last10-Middle]] | Main | 0.073 | -0.009 – 0.156 | 4139.128 | 1.736 | 0.0826 |
| Sex[H.Male-Female] ×<br>Position[H.First10-avg(Last10,Middle)] | Main | 0.019 | -0.017 – 0.054 | 3722.516 | 1.039 | 0.2991 |
| Solution[H.Sucrose-Water] ×<br>Position[H.Last10-Middle]] | Main | 0.002 | -0.094 – 0.098 | 3678.942 | 0.038 | 0.9693 |
| Solution[H.EtOH-avg(S,W)] ×<br>Position[H.Last10-Middle]] | Main | 0.013 | -0.048 – 0.074 | 3464.364 | 0.416 | 0.6776 |
| Solution[H.Sucrose-Water] ×<br>Position[H.First10-avg(Last10,Middle)] | Main | 0.064 | 0.024 – 0.103 | 4404.965 | 3.119 | **0.0018** |
| Solution[H.EtOH-avg(S,W)] ×<br>Position[H.First10-avg(Last10,Middle)] | Main | -0.020 | -0.044 – 0.005 | 4167.059 | -1.548 | 0.1217 |
| time[H.After-Before] ×<br>Sex[H.Male-Female] ×<br>Solution[H.Sucrose-Water] | Main | 0.003 | -0.053 – 0.060 | 3526.440 | 0.121 | 0.9036 |
| time[H.After-Before] ×<br>Sex[H.Male-Female] ×<br>Solution[H.EtOH-avg(S,W)] | Main | -0.022 | -0.055 – 0.012 | 3554.010 | -1.273 | 0.2033 |
| time[H.After-Before] ×<br>Sex[H.Male-Female] ×<br>Position[H.Last10-Middle]] | Main | -0.016 | -0.080 – 0.049 | 3548.776 | -0.484 | 0.6285 |
| time[H.After-Before] ×<br>Sex[H.Male-Female] ×<br>Position[H.First10-avg(Last10,Middle)] | Main | 0.003 | -0.024 – 0.029 | 3554.170 | 0.183 | 0.8552 |
| time[H.After-Before] ×<br>Solution[H.Sucrose-Water] ×<br>Position[H.Last10-Middle]] | Main | 0.004 | -0.073 – 0.082 | 3553.468 | 0.114 | 0.9091 |
| time[H.After-Before] ×<br>Solution[H.EtOH-avg(S,W)] ×<br>Position[H.Last10-Middle]] | Main | -0.016 | -0.063 – 0.030 | 3540.658 | -0.688 | 0.4916 |
| time[H.After-Before] ×<br>Solution[H.Sucrose-Water] ×<br>Position[H.First10-avg(Last10,Middle)] | Main | -0.008 | -0.041 – 0.025 | 3553.866 | -0.480 | 0.6312 |
| time[H.After-Before] ×<br>Solution[H.EtOH-avg(S,W)] ×<br>Position[H.First10-avg(Last10,Middle)] | Main | 0.011 | -0.009 – 0.030 | 3553.593 | 1.090 | 0.2758 |
| Sex[H.Male-Female] ×<br>Solution[H.Sucrose-Water]<br>× Position[H.Last10-Middle]] | Main | 0.011 | -0.085 – 0.107 | 3678.496 | 0.226 | 0.8214 |
| Sex[H.Male-Female] ×<br>Solution[H.EtOH-avg(S,W)] ×<br>Position[H.Last10-Middle]] | Main | 0.011 | -0.051 – 0.072 | 3468.730 | 0.342 | 0.7326 |
| Sex[H.Male-Female] ×<br>Solution[H.Sucrose-Water]<br>× Position[H.First10-avg(Last10,Middle)] | Main | 0.027 | -0.013 – 0.067 | 4386.700 | 1.334 | 0.1823 |
| Sex[H.Male-Female] ×<br>Solution[H.EtOH-avg(S,W)]<br>× Position[H.First10-avg(Last10,Middle)] | Main | -0.010 | -0.035 – 0.015 | 4169.119 | -0.808 | 0.4193 |
| time[H.After-Before] ×<br>Sex[H.Male-Female] ×<br>Solution[H.Sucrose-Water] ×<br>Position[H.Last10-Middle]] | Main | -0.015 | -0.092 – 0.062 | 3553.455 | -0.388 | 0.6983 |
| time[H.After-Before] ×<br>Sex[H.Male-Female] ×<br>Solution[H.EtOH-avg(S,W)] ×<br>Position[H.Last10-Middle]] | Main | -0.018 | -0.065 – 0.028 | 3540.652 | -0.761 | 0.4466 |
| time[H.After-Before] ×<br>Sex[H.Male-Female] ×<br>Solution[H.Sucrose-Water] ×<br>Position[H.First10-avg(Last10,Middle)] | Main | 0.007 | -0.026 – 0.040 | 3553.839 | 0.428 | 0.6685 |
| time[H.After-Before] ×<br>Sex[H.Male-Female] ×<br>Solution[H.EtOH-avg(S,W)] ×<br>Position[H.First10-avg(Last10,Middle)] | Main | -0.002 | -0.021 – 0.017 | 3553.597 | -0.190 | 0.8497 |
| time[After-Before] ×<br>Solution[EtOH]] ×<br>Position[First10-Last10]] | Post-Hoc | -0.0241 | -0.509 – 0.460 | 3818.728 | -0.098 | 0.9222 |
| time[After-Before] ×<br>Solution[Water]] ×<br>Position[First10-Last10]] | Post-Hoc | -0.257 | -0.841 – 0.327 | 3822.387 | -0.863 | 0.3882 |
| time[After-Before] ×<br>Solution[Sucrose] ×<br>Position[First10-Last10] | Post-Hoc | -0.371 | -0.671 – -0.072 | 3854.694 | -2.428 | **0.0152** |
| **Figure 5 – CeA<sup>Dyn</sup> Calcium Transients in Response to Negative Valence, Valuation State, and Taste** |  |  |  |  |  |  |
| **Figure 5A & B – Consumption (mL/kg)** |  |  |  |  |  |  |
| (Intercept) | Main | 31.572 | 22.847 – 40.298 | 6.198 | 8.786 | **0.0001** |
| Sex[E.Male-Female] | Main | -3.850 | -12.575 – 4.876 | 6.198 | -1.071 | 0.3240 |
| Solution[E.QHCl-Water] | Main | -4.293 | -15.475 – 6.889 | 36.428 | -0.778 | 0.4414 |
| Solution[E.WaterDep-Water] | Main | 50.240 | 37.062 – 63.419 | 36.105 | 7.731 | **<0.0001** |
| Solution[E.S+QHCl<sub>low</sub>-Water] | Main | 2.504 | -9.955 – 14.962 | 36.411 | 0.407 | 0.6861 |
| Solution[E.S+QHCl<sub>high</sub>-Water] | Main | 2.401 | -11.269 – 16.071 | 36.820 | 0.356 | 0.7239 |
| Sex[E.Male-Female] ×<br>Solution[E.QHCl-Water] | Main | -1.314 | -12.496 – 9.868 | 36.428 | -0.238 | 0.8131 |
| Sex[E.Male-Female] ×<br>Solution[E.WaterDep-Water] | Main | -10.532 | -23.711 – 2.646 | 36.105 | -1.621 | 0.1138 |
| Sex[E.Male-Female] ×<br>Solution[E.S+QHCl<sub>low</sub>-Water] | Main | -0.991 | -13.450 – 11.468 | 36.411 | -0.161 | 0.8728 |
| Sex[E.Male-Female] ×<br>Solution[E.S+QHCl<sub>high</sub>-Water] | Main | -8.833 | -22.503 – 4.837 | 36.820 | -1.309 | 0.1985 |
| Solution[S+QHCl<sub>high</sub>-<br>S+QHCl<sub>low</sub>] | Post-Hoc | -0.1028 | -19.205 – 18.999 | 37.706 | -0.015 | 1 |
| **Figure 5C & E – GCaMP7f Signal (z%ΔF/F)** |  |  |  |  |  |  |
| (Intercept) | Main | 0.008 | -0.198 – 0.213 | 16.793 | 0.079 | 0.9383 |
| time[E.After-Before] | Main | 0.151 | 0.093 – 0.208 | 14.348 | 5.610 | **0.0001** |
| Solution[E.QHCl-Water] | Main | -0.654 | -0.811 – -0.496 | 1064.402 | -8.147 | **<0.0001** |
| Solution[H.WDep-Water] | Main | 0.322 | 0.207 – 0.436 | 2070.154 | 5.506 | **<0.0001** |
| Solution[E.S+QHCl<sub>low</sub>-Water] | Main | -0.087 | -0.208 – 0.033 | 2079.855 | -1.424 | 0.1546 |
| Solution[E.S+QHCl<sub>high</sub>-Water] | Main | -0.173 | -0.310 – -0.037 | 2061.311 | -2.488 | **0.0129** |
| Sex[E.Male-Female] | Main | -0.115 | -0.318 – 0.087 | 15.744 | -1.210 | 0.2442 |
| time[E.After-Before] ×<br>Solution[E.QHCl-Water] | Main | -0.075 | -0.190 – 0.041 | 162.634 | -1.280 | 0.2023 |
| time[E.After-Before] ×<br>Solution[E.WaterDep-Water] | Main | -0.014 | -0.118 – 0.090 | 1168.597 | -0.262 | 0.7935 |

| **Contrast**<br>Factor [Contrast] x Interaction | **Type** | **Estimate** | **95% CI** | **df** | **Statistic** | **p Value** |
| --- | --- | --- | --- | --- | --- | --- |
| time[E.After-Before] ×<br>Solution[E.S+QHCl low-Water] | Main | -0.062 | -0.174 – 0.050 | 1544.703 | -1.086 | 0.2777 |
| time[E.After-Before] ×<br>Solution[E.S+QHCl high-Water] | Main | -0.097 | -0.220 – 0.027 | 1309.918 | -1.528 | 0.1268 |
| time[E.After-Before] ×<br>Sex[E.Male-Female] | Main | 0.055 | -0.002 – 0.113 | 14.349 | 2.059 | 0.0581 |
| Solution[E.QHCl-Water] ×<br>Sex[E.Male-Female] | Main | 0.085 | -0.072 – 0.243 | 1053.275 | 1.061 | 0.2890 |
| Solution[E.WaterDep-Water] ×<br>Sex[E.Male-Female] | Main | 0.018 | - 0.097 – 0.132 | 2071.353 | 0.300 | 0.7638 |
| Solution[E.S+QHCl low-Water] ×<br>Sex[E.Male-Female] | Main | 0.279 | 0.159 – 0.400 | 2082.233 | 4.549 | **<0.0001** |
| Solution[E.S+QHCl high-Water]<br>×<br>Sex[E.Male-Female] | Main | 0.109 | -0.027 – 0.246 | 2062.054 | 1.570 | 0.1167 |
| time[E.After-Before] ×<br>Solution[E.QHCl-Water] ×<br>Sex[E.Male-Female] | Main | 0.022 | -0.093 – 0.137 | 162.634 | 0.377 | 0.7070 |
| time[E.After-Before] ×<br>Solution[E.WaterDep-Water] ×<br>Sex[E.Male-Female] | Main | 0.056 | -0.047 – 0.160 | 1168.610 | 1.065 | 0.2869 |
| time[E.After-Before] ×<br>Solution[E.S+QHCl low-Water] ×<br>Sex[E.Male-Female] | Main | 0.125 | 0.013 – 0.238 | 1544.707 | 2.192 | **0.0285** |
| time[E.After-Before] ×<br>Solution[E.S+QHCl high-Water] ×<br>Sex[E.Male-Female] | Main | 0.016 | -0.108 – 0.140 | 1309.922 | 0.254 | 0.7995 |
| time[After-Before] ×<br>Solution[Water] ×<br>Sex[Female-Male] | Post-Hoc | -0.397 | -0.771 – -0.023 | 79.854 | -2.112 | **0.0378** |
| Solution[S+QHCl low – Water] ×<br>Sex[Female] | Post-hoc | -0.367 | -0.591 – -0.142 | 2116.302 | -3.830 | **0.0004** |
| Solution[S+QHCl low – Water] ×<br>Sex[Male] | Post-hoc | 0.192 | 0.011 – 0.372 | 1964.290 | 2.495 | **0.0339** |
| time[After-Before] ×<br>Solution[S+QHCl low – Water] ×<br>Sex[Female] | Post-hoc | -0.375 | -0.715 – -0.035 | 1259.405 | -2.162 | **0.0308** |
| time[After-Before] ×<br>Solution[S+QHCl low – Water] ×<br>Sex[Male] | Post-hoc | 0.127 | -0.166 – 0.419 | 1780.402 | 0.848 | 0.3963 |
| Solution[E.QHCl] × Sex[E.Male-Female] | Post-hoc | -0.068 | -0.496 – 0.359 | 20.707 | 0.332 | 0.7431 |
| **Figure S1 – CeA<sup>dyn</sup> Calcium Transients in the First Bout of the First Session** |  |  |  |  |  |  |
| **Figure S1B – First Bout GCaMP7f Signal (z%ΔF/F)** |  |  |  |  |  |  |
| (Intercept) | Main | 0.018 | -1.310 – 1.347 | 3.00 | 0.044 | 0.9675 |
| time[H.After-Before] | Main | 0.520 | 0.140 – 0.901 | 15.00 | 2.918 | **0.0106** |
| Solution[H.Sucrose-Water] | Main | -0.137 | -0.602 – 0.329 | 15.00 | -0.625 | 0.5414 |
| Solution[H.EtOH-avg(S,W)] | Main | 0.242 | -0.027 – 0.511 | 15.00 | 1.918 | 0.0744 |
| time[H.After-Before] ×<br>Solution[H.Sucrose-Water] | Main | 0.001 | -0.464 – 0.467 | 15.00 | 0.006 | 0.9950 |
| time[H.After-Before] ×<br>Solution[H.EtOH-avg(S,W)] | Main | 0.306 | 0.037 – 0.575 | 15.00 | 2.425 | **0.0284** |
| time[H.After-Before] ×<br>Solution[EtOH] | Post-hoc | 2.265 | 0.947 – 3.582 | 15.00 | 3.665 | **0.0023** |
| time[H.After-Before] ×<br>Solution[Water] | Post-hoc | 0.426 | -0.891 – 1.743 | 15.00 | 0.690 | 0.5010 |
| time[H.After-Before] ×<br>Solution[Sucrose] | Post-hoc | 0.432 | -0.885 – 1.749 | 15.00 | 0.699 | 0.4950 |

## Supplement C – Supplemental Data

**Figure S1.**
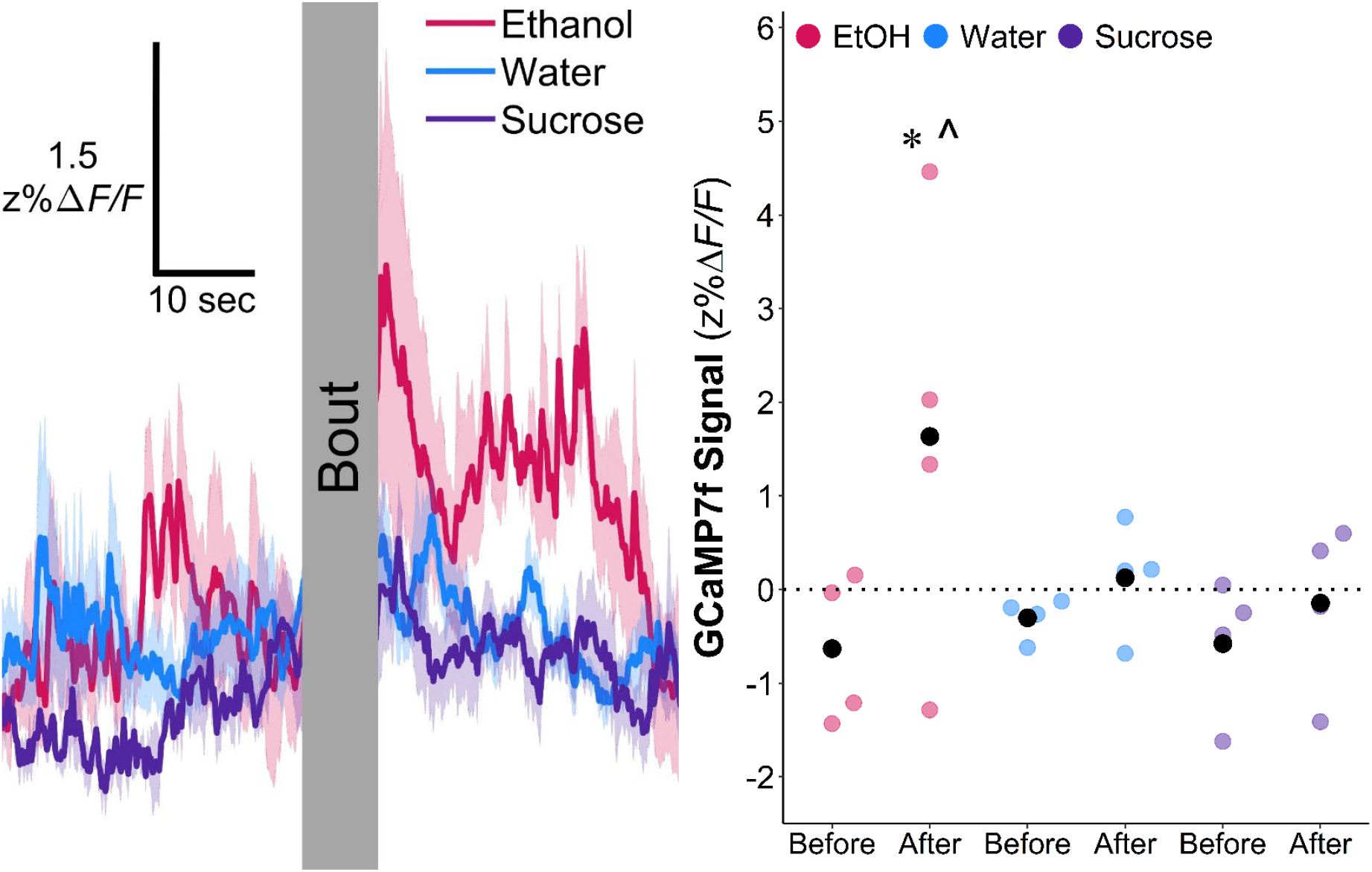
Calcium activity during the first drinking bout. Since not all animals were recorded during their first exposure, the traces include data from four animals, two of each sex, around the first EtOH and sucrose bouts within the first exposure sessions for each solution. Water, though previously experienced, was included for comparison. (**A**) Mean ± SE GCaMP7f signal (z%ΔF/F) traces 30 seconds before and 30 seconds after drinking bouts and (**B**) average GCaMP7f signal (z%ΔF/F) 1 sec before and after individual drinking bouts, both collapsed across sex. Despite never having experienced intoxication or other ingestive effects of EtOH, the increase in calcium signal associated with an EtOH bout is markedly more pronounced than the average sucrose and water signals, indicating that the response described in the study is likely innate. Signal MLMM with subject random intercept; *, significantly higher than respective before-bout signal, p <.05; ^, significantly higher signal increase around bout compared to water and sucrose, p <.05.

## Notes

### Competing Interest Statement

The authors have declared no competing interest.

### Summary of Updates

The revised manuscript contains refined and new analysis.

